# ARTIFICIAL RECOGNITION ELEMENTS FOR AFRICAN SWINE FEVER VIRUS: DNA APTAMERS WITH UNUSUAL NI^2+^-BINDING MOTIFS

**DOI:** 10.64898/2026.05.05.722837

**Authors:** Rugiya Aliyeva, Vladimir Mushenkov, Hovakim Zakaryan, Astghik Hakobyan, Tigran Harutyunyan, Galina Hovhannisyan, Anzhela Sargsyan, Nadezda Meshcheryakova, Olga Zaborova, Ilya Oleynikov, Uliana Bulgakova, Sofia Mariasina, Liliya Mukhametova, Sergei Eremin, Galina Koltsova, Rouben Aroutiounian, Yulia Gorshkova, Alexander Nechaev, Vladimir Tsvetkov, Elena Zavyalova

## Abstract

Rapid and accurate diagnosis of viral and bacterial infections is a significant challenge in medicine and veterinary science, especially in the case of epidemically dangerous pathogens. The African swine fever virus (ASFV) causes annual outbreaks among livestock, causing in significant economic losses for farmers. Artificial recognition elements, DNA aptamers, have been identified as a suitable tool for point-of-care diagnostics due to their high affinity and specificity for the target, as well as stability at ambient temperatures during storage. This study reports for the first time on DNA aptamers that can bind ASFV particles. The aptamers were generated using a single-round selection process; they bound to both recombinant p54 protein and ASFV particles as well as exhibited antiviral properties. Functional activity of one of the aptamers depended on Ni^2+^, which was a dopant during the selection process; this cation increased aptamer affinity by 100-fold. A combination of techniques was used to predict the structure of Ni^2+^-regulated aptamer identifying a new non-canonical DNA motif. Cation-dependent regulation of aptamer activity is promising as it allows regeneration of affine carriers and biosensors with chelating agents, while addition of Ni^2+^ restores the high-affine aptamer conformation.

## Introduction

African swine fever is a highly contagious and devastating viral disease that affects domestic pigs and wild boar, causing significant economic losses [1,2]. African swine fever virus (ASFV) is a large double-stranded DNA virus that belongs to the *Asfarviridae* family [3]. It was first identified in Kenya in 1921 and has since spread to other parts of the world, including Europe and Asia [4,5]. The lack of a commercially available vaccine for the virus makes it difficult to control the disease, and early detection is essential to prevent its spread [6].

The ASFV genome encodes more than 150 proteins [7]. Among these proteins, the structural protein p54 has been identified as a promising candidate for diagnostic development. p54 is a homodimeric transmembrane protein located in the viral envelope and is essential for the viability of the virus. Repression of p54 synthesis prevents the formation of viral envelope membranes and arrests virus morphogenesis at an early stage [8]. p54 also plays a role in the early stages of infection by interacting with host cell dynein motors, facilitating the transport of the virus within the cell [9]. Both purified p54 protein and antibodies against p54 have been shown to prevent infection by inhibiting the attachment of viral particles to host cells [10]. During the infection, anti-p54 antibodies are detected as early as 8 days after infection and persist for several weeks; they are an attractive target for diagnosis [11,12].

Given its strong immunogenicity and early expression, p54 has been widely validated as a promising biomarker. Enzyme-linked immunosorbent assays (ELISA) showed 98% sensitivity and 97% specificity in detecting antibodies to ASFV [13,14]. To facilitate point-of-care testing, lateral flow immunochromatographic assays (LFIA) were developed to detect ASFV antibodies. Although colloidal gold-based LFIA had a nearly identical limit of detection to ELISA, quantum dot-based LFIA offered improved performance with a 1000-fold lower detection limit [15, 16]. Despite the widespread use of antibody detection, the delayed appearance of antibodies *in vivo* can lead to false-negative tests in the early stages of infection. Detection of the virus itself could address this issue. The polymerase chain reaction (PCR) remains the most accurate method [17], but it requires specialized laboratory equipment and skilled personnel to avoid false positive results caused by sample contamination.

Nucleic acid aptamers are a class of artificial recognition molecules with high affinity and specificity. These molecules are a synthetic alternative to antibodies having very broad range of possible targets. The aptamers are selected from combinatorial libraries through a process called systematic evolution of ligands by exponential enrichment (SELEX). Chemical synthesis of aptamers allows for site-specific modifications with various functional groups, making them suitable for a wide range of applications [18,19]. Dozens of aptamers have selected for different viruses [20], but aptamers specific to ASFV have not yet been described. A recent study [21] described DNA aptamers for p30, a protein expressed by ASFV-infected cells. p30 is not exposed on the surface of the virus, but it can be found in serum samples from infected animals. The aptamers showed a high affinity for the target with dissociation constants (*K*_*D*_) ranging from 140 to 270 pM. An enzyme-linked aptamer assay for the p30 protein achieved a detection limit of 0.61 ng/mL, which is achievable during the acute phase of ASF.

At the moment of the first preprint of this study, no specific aptamer has been reported for the surface epitopes of ASFV. Given the diagnostic value of p54 and the inherent advantages of aptamers as recognition molecules, the selection of a p54-specific aptamer is a logical and important next step. In this study, we describe successful *in vitro* selection of a DNA aptamer that targets the recombinant ASFV p54 protein using a single-round selection procedure. The aptamers have no sequence similarity to reported recently aptamers to the same protein [22]. For the first time, aptamers to p54 have demonstrated the ability to bind virions and to decrease viral titer during infection. Additionally, we have identified an unusual DNA motif that binds to Ni^2+^. This motif may have a practical role as cation-dependent regulation of aptamer activity allows regeneration of affine carriers and biosensors with chelating agents, while addition of Ni^2+^ can restore the high-affine aptamer conformation.

## Results and Discussion

### DESIGN OF THE LIBRARY AND APTAMER SELECTION THROUGH A SINGLE ROUND

The library of 58-nucleotide DNA oligonucleotides consisted of 3 regions with randomized sequences (12, 6, and 12 nucleotides), as well as a hairpin stabilized by 6 pairs of nucleotides, forming an element of the secondary DNA structure. The introduction of a structural element was shown to be an effective approach for aptamer selection [23,24]. 8 non-randomized nucleotides were added at the 5’- and 3’-ends of the sequence in order to allow for PCR amplification and sequencing of the library (Supplementary figure 1). The total number of randomized nucleotides was 30, corresponding to 10^18^possible variants of oligonucleotides. A theoretical library containing all possible variants would have a size of 1.9 µmoles or 37 mg of oligonucleotides. For the selection experiment, 740 nmoles of oligonucleotides were synthesized, representing approx. 39% of theoretical maximum library size.

**Figure 1.**
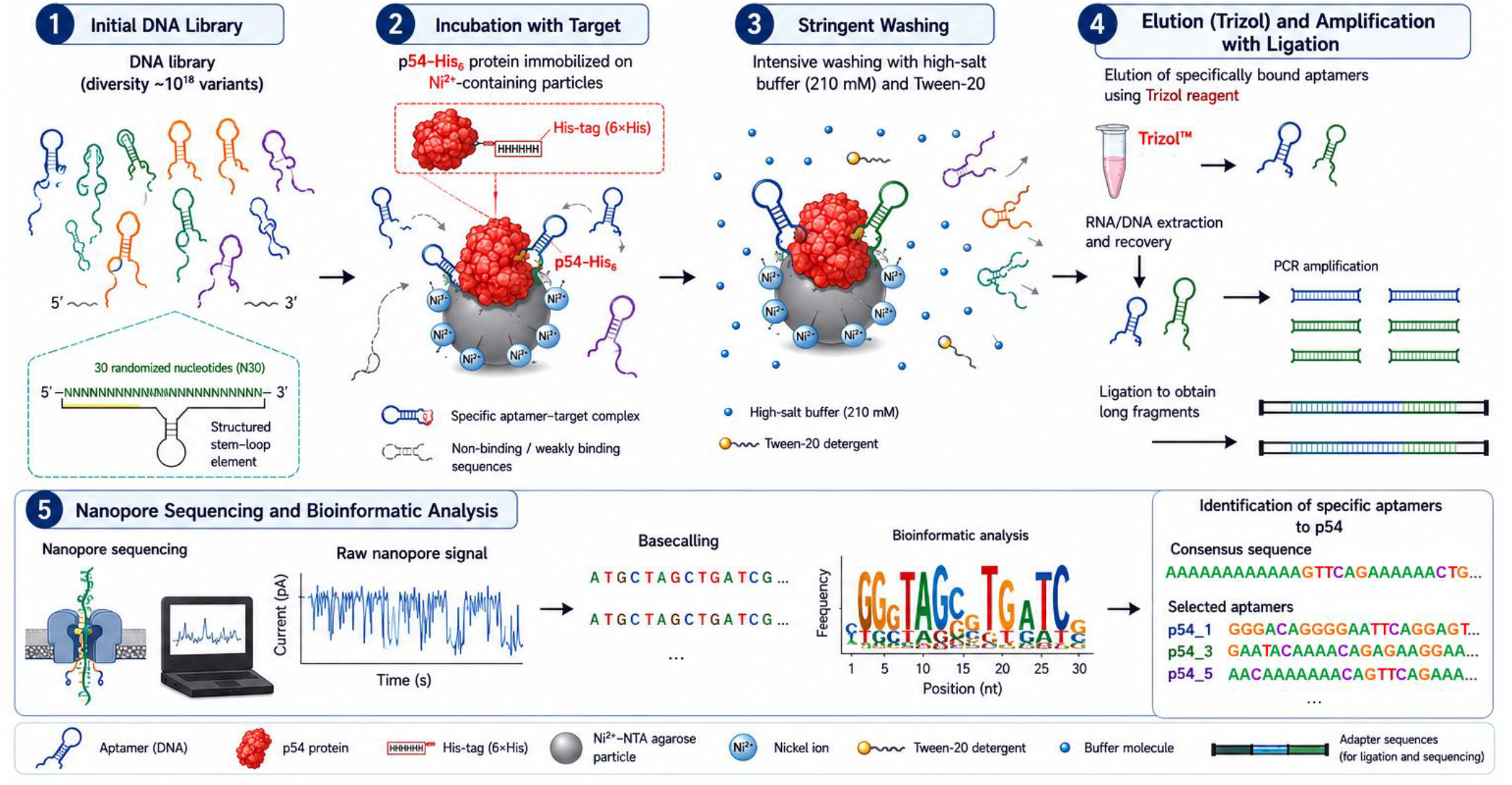
A schematic representation of the single-round selection process for DNA aptamers against recombinant p54 protein. The icons for this figure were generated using ChatGPT and arranged manually.

Aptamer selection is a multistep process that involves repetitive rounds of selecting oligonucleotides with high affinity and amplifying them. In addition to traditional approaches, single-round aptamer selection was described in several studies. These include selecting aptamers with high affinity by eliminating low-specific sequences through intensive washing (MonoLEX) [25,26] or hydrolyzing them with nucleases [27]. We followed the MonoLEX approach, but modified it technically, while maintaining the general idea of the selection process. Ni^2+^-containing particles were coated with p54 protein that bears a His_6_-tag. After interacting with a DNA library, the non-affine sequences were washed out with a 900-fold excess of the buffer containing an increasing amount of a Tween-20 detergent to decrease hydrophobic-driven adsorption. The selection buffer had an excessive ionic strength of 210 mM, which reduces the electrostatic-driven adsorption. After a single round of selection, the aptamer-rich library was sequenced (Figure 1).

Nanopore sequencing was chosen to identify the composition of the aptamer-enriched library as it can provide good coverage. This type of sequencing was used in aptamer selection previously [28], but it required crosslinking of aptamers to obtain elongated DNA fragments. In this study, we attempted to elongate DNA using complementary primers as well as T4 DNA ligase. Additionally, the DNA length was increased with a barcoding kit. Despite these approaches, most sequenced fragments correspond to the unligated PCR product (approx. 195 base pairs, including barcoding sequences), with minor fractions of dimeric and trimeric variants (Supplementary figure 2). Generally, short fragments are not optimal for nanopore cell operation, but the quality of the reads was sufficient for these short fragments. Thus, nanopore sequencing with 10^th^ generation chemistry combined with barcoding allowed for the direct acquisition of aptamer sequences after PCR amplification, without the need for additional processing steps.

**Figure 2.**
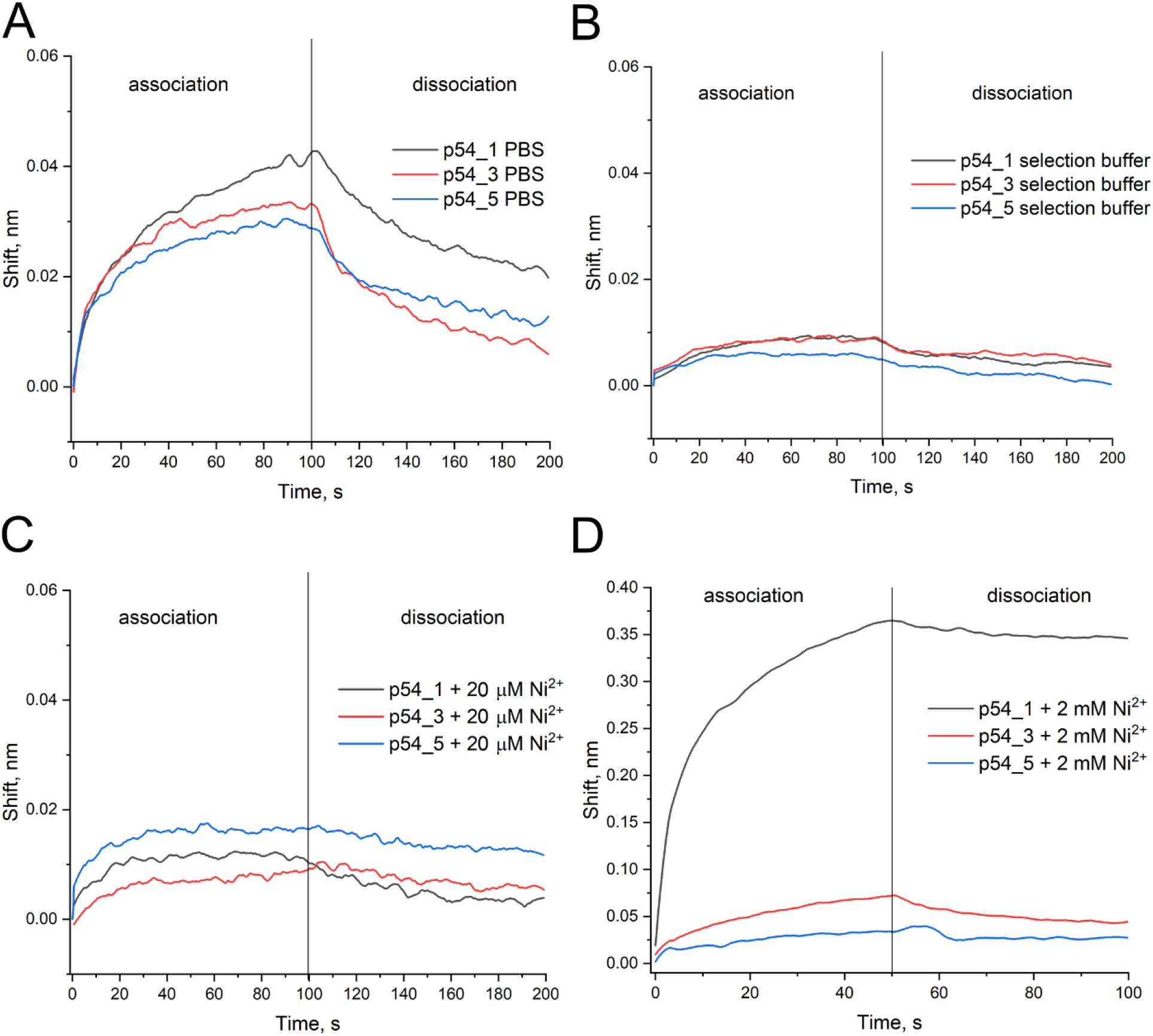
Affinity estimation of aptamers for p54 protein using biolayer interferometry. The complex association and dissociation in PBS (a), selection buffer (b), selection buffer with 20 µM NiCl_2_ (c), and selection buffer with 2 mM NiCl_2_ (d), for three aptamers (p54_1, p54_3, p54_5). The curves were acquired for 1 µM solutions of each aptamer.

An analysis of the results revealed the presence of 3,380 DNA sequences that matched the library’s pattern. Of these, 99.5% were unique. Bioinformatic analysis identified a global consensus sequence with randomized regions enriched with adenines (a probability of about 60%, Supplementary table 1). Cytosines were the second most common nucleotide in the first and second randomized regions, with a probability of approx. 15%. In the third randomized region, other nucleotides had similar probabilities (10-13%). In addition to the global consensus sequence, several local consensus sequences were selected for further investigation (Supplementary table 2).

**Table 1.** *K*_*D*_ estimation of aptamer-p54 protein complexes in different buffers. Values are given in nM.

| Aptamer | PBS | Selection buffer | Selection buffer with 20 $\mu\text{M}$ $\text{Ni}^{2+}$ | Selection buffer with 2 mM $\text{Ni}^{2+}$ |
| --- | --- | --- | --- | --- |
| p54_1 | 54±9 | 400±90 | 37±19 | 4±2 |
| p54_3 | 30±11 | 42±18 | 60±20 | 15±5 |
| p54_5 | 75±19 | 170±40 | 110±30 | 160±30 |

**Table 2.** *K*_*D*_ estimation for complexes of modified aptamers with p54 protein in selection buffer containing 20 µM Ni^2+^. Values are given in nM. Alexa488 is Alexa dye with an excitation wavelength of 488 nm, and FAM is carboxyfluorescein.

| Aptamer | No modification | Alexa488 | FAM | Biotin |
| --- | --- | --- | --- | --- |
| PBS |  |  |  |  |
| p54_1 | 54±9 | 50±17 | 37±8 | 30±9 |
| p54_3 | 30±11 | 15±5 | 29±7 | 21±9 |
| p54_5 | 75±19 | 34±16 | 80±30 | 90±30 |
| Selection buffer |  |  |  |  |
| p54_1 | 400±90 | 760±130 | 115±13 | 580±110 |
| p54_3 | 42±18 | 140±40 | 38±3 | 62±18 |
| p54_5 | 170±40 | 31±15 | 39±9 | 800±300 |
| Selection buffer with 20 $\mu\text{M}$ $\text{Ni}^{2+}$ | | | | |
| p54_1 | 37±19 | n.d. | 23±5 | 88±12 |
| p54_3 | 60±20 | 32±15 | 60±20 | 90±30 |
| p54_5 | 110±30 | 39±12 | 70±20 | 80±30 |

### AFFINITY OF THE APTAMERS IS HIGHLY DEPENDENT ON THE BUFFER USED AND THE LABELS INTRODUCED

The affinity of the global consensus sequence and 10 local consensus sequences for immobilized p54 protein was examined using biolayer interferometry (Figure 2, Table 1). Among all variants studied, only three variants showed binding curves (Supplementary figure 3), namely p54_1, p54_3, and p54_5. Notably, the global consensus sequence did not show notable affinity for p54. The affinities of 2 out of 3 aptamers were depended on the buffer system used. p54_1 had *K*_*D*_ of 54 nM in PBS and 400 nM in the selection buffer. Next, we hypothesized that other dopants might also affect the performance of the aptamers. We tested Ni^2+^ ions, which are a component of the particles for protein immobilization. The addition of 20 µM Ni^2+^ increased the affinity of p54_1 by an order of the magnitude, yielding *K*_*D*_ of 37 nM. While 2 mM Ni^2+^ further increased the affinity, yielding *K*_*D*_ of 4 nM. The effects on p54_3 and p54_5 were less significant, with 20 µM Ni^2+^ slightly increasing the bound aptamer fraction of p54_5 and having a minor effect on the *K*_*D*_ value.

**Figure 3.**
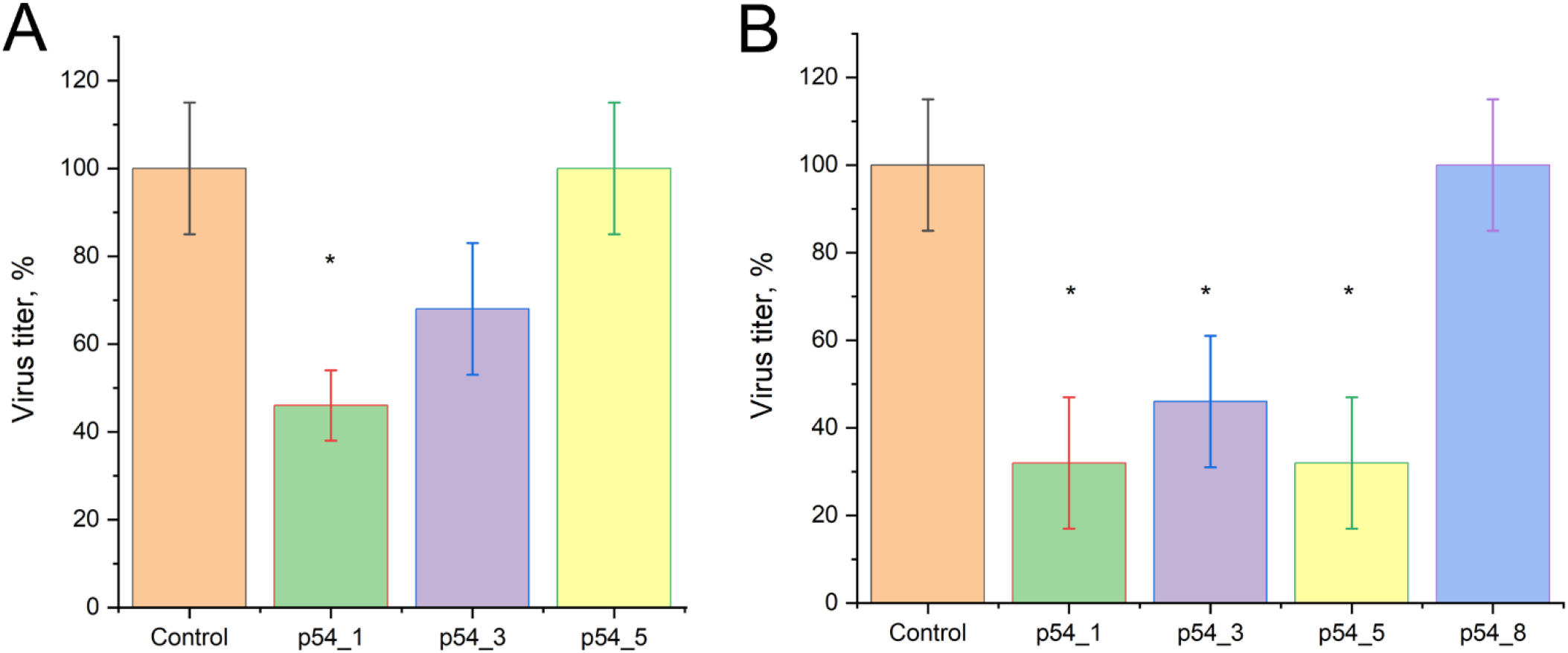
The antiviral effects of DNA aptamers against p54 estimated in ASFV infection of Vero cell cultures. Co-treatment of the cells with 10 µM aptamers and BA71V strain of ASFV; the effect was estimated 72 hours after infection (a). Sequential treatment of the cells with BA71V strain of ASFV and then, 16 hours later, with 10 µM of aptamers; the antiviral effect was estimated 120 hours after infection (b). Asterisks indicate statistically significant differences with p-values less than 0.05.

Next, we decided to estimate the affinity of the aptamers for viruses. All current methods for virus binding require aptamer modification. The effect of aptamer modification was investigated in PBS, selection buffer, and selection buffer containing 20 µM Ni^2+^ (Table 2). The affinity for the recombinant protein varied depending on the modification. The most significant effects were observed for Alexa488 derivatives, namely, p54_1_AF showed low affinity for the protein in the selection buffer, while p54_3_AF and p54_5_AF affinities increased by 2-3 fold. FAM modification also increased affinity, while biotin modification decreased affinity in selection buffer. Several papers described approaches for aptamer affinity estimation [29,30], discussing the possible effect of aptamer labeling on affinity. Our data are consistent with these discussions. For the further experiments with viruses, PBS was chosen, as it provided the highest averaged affinity with minor effects of modifications.

### DNA APTAMERS BIND ASFV VIRIONS

The protein p54 is a membrane dimeric protein [31], and its interaction with recognition molecules can be challenging due to steric constraints [32]. The binding of recombinant p54 protein does not necessarily guarantee aptamer binding to viral particles. To address this issue, we tested the interaction between aptamers and viral particles using two strains: Stavropol_01/08 (Genotype II, Serogroup 8, GenBank accession number PQ672299.1) [33] and KK262 (Congo-a) (Genotype I, Serogroup 2, GenBank accession number OM249788.1) [34].

Biolayer interferometry was performed using biotinylated aptamers immobilized on a sensor. The KK262 virus bound to all three aptamers in PBS with a *K*_*D*_ of 7-10·10^7^ viral particles/mL (Supplementary figure 4, Table 3). However, only p54_3 bound to the Stavropol_01/08 virus with a *K*_*D*_ of 6·10^5^ viral particles/mL, indicating an increase in affinity by 2 orders of magnitude compared to the KK262 strain.

**Table 3.** *K*_*D*_ estimation for aptamer complexes with ASFV virions in PBS. The values are given in viral particles/mL. n.d. is no binding. The values for nanoparticle tracking analysis represent fractions of virions labeled with aptamers.

| Aptamer | Biolayer interferometry |  | Fluorescence polarization |  | Nanoparticle tracking analysis |  |
| --- | --- | --- | --- | --- | --- | --- |
|  | KK262 | Stavropol_01/08 | KK262 | Stavropol_01/08 | KK262 | Stavropol_01/08 |
| p54_1 | $(7.0 \pm 1.1) \cdot 10^7$ | n.d. | $(8 \pm 2) \cdot 10^7$ | n.d. | 4% | 4% |
| p54_3 | $(10 \pm 4) \cdot 10^7$ | $(6 \pm 3) \cdot 10^5$ | n.d. | $(1.4 \pm 0.4) \cdot 10^8$ | 8% | 3% |
| p54_5 | $(9 \pm 3) \cdot 10^7$ | n.d. | $(2.2 \pm 0.9) \cdot 10^8$ | n.d. | 20% | 9% |

**Figure 4.**
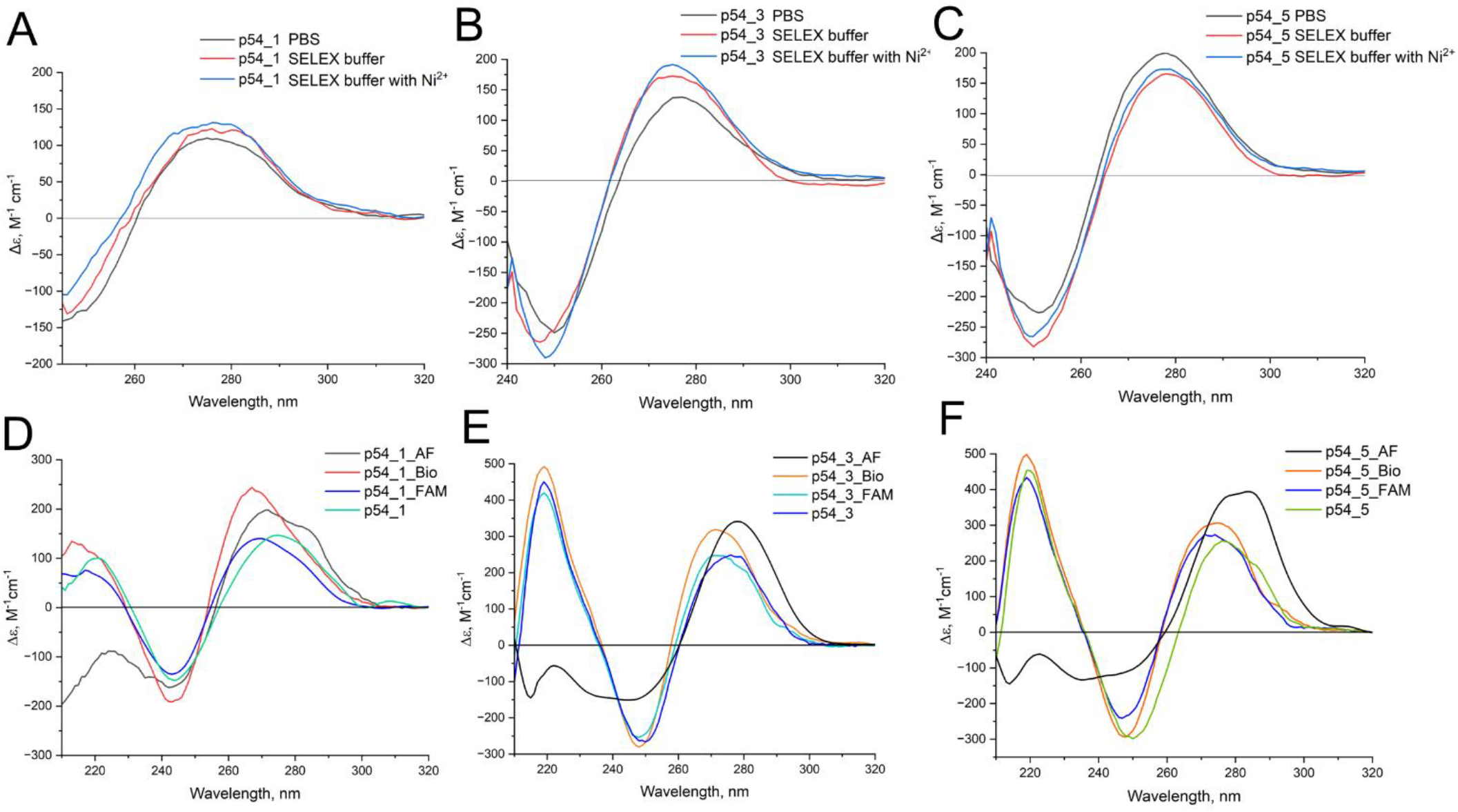
Circular dichroism spectra of p54_1 (A), p54_3 (B), and p54_5 (C) in different buffers. The effect of aptamer modification on their circular dichroism spectra in the selection buffer containing 20 µM NiCl_2_: p54_1 (D), p54_3 (E), and p54_5 (F). The curves were obtained for a 1 µM solution of each aptamer.

The binding of aptamers to viruses was also assessed in homogeneous systems. For the fluorescence polarization assay, the aptamers were FAM-modified. The addition of virus increased the polarization of fluorescence by 10-20%, due to a slowing down of the rotation of the fluorescent label. Binding of p54_1 and p54_5 to ASFV KK262 was observed, as well as p54_3 bound to Stavropol_01/08 (Supplementary figure 5, Table 3). Dissociation constants for p54_1 determined by different methods overlapped within the error margin. For p54_3, there was a significant difference in affinity of 200-fold, which may be due to the conformational change caused by modification.

**Figure 5.**
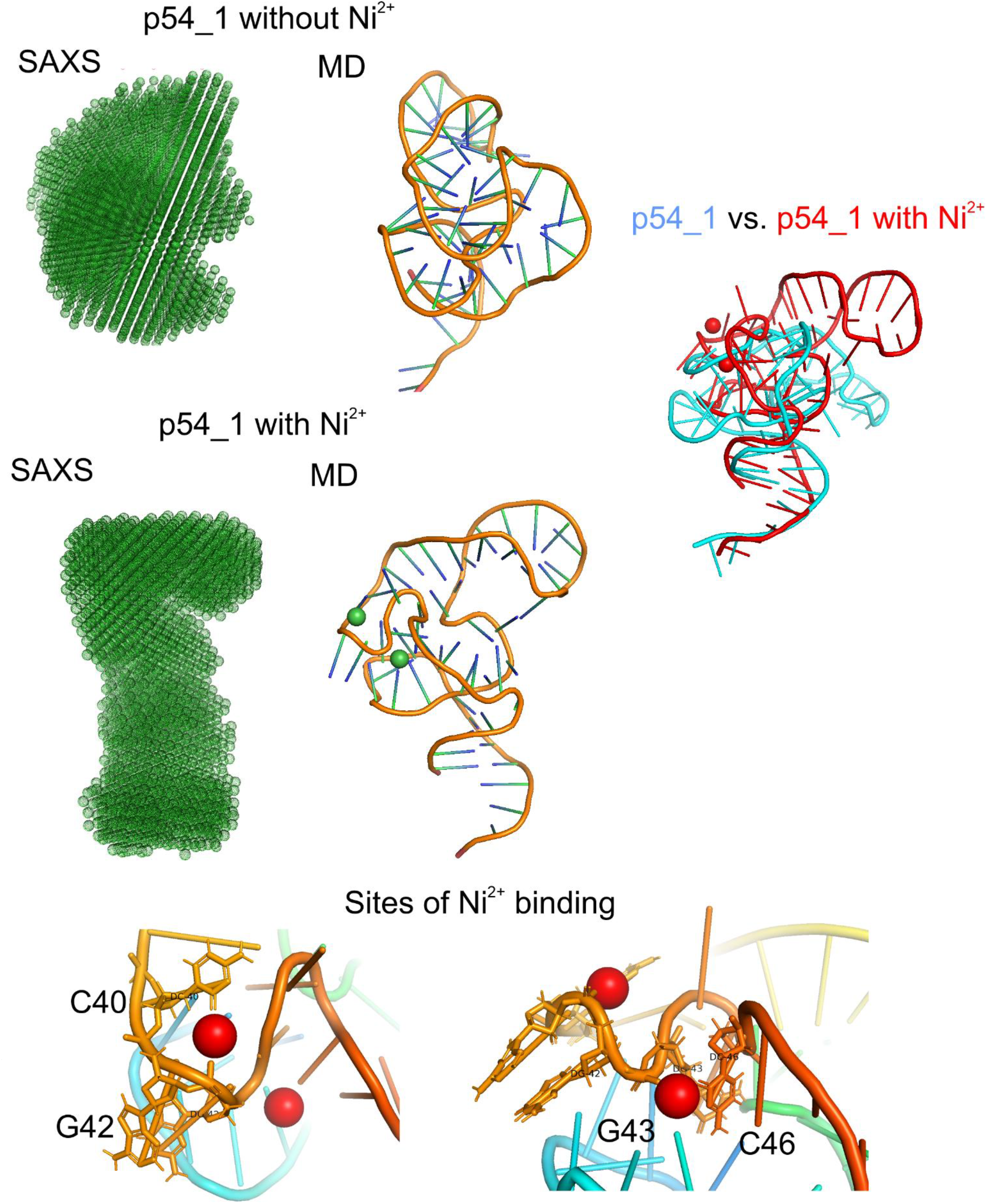
Proposed structures of p54_1 and its complex with 2 Ni^2+^ ions. SAXS determines aptamer conformation, which is switched in the presence of Ni^2+^ ions. Molecular dynamics simulations were used to propose the conformations, taking into account NMR data. Each Ni^2+^ ion is coordinated with one phosphate group and oxygen of cytosine that caused distortion of the aptamer backbone rearranging loops into a new form.

Nanoparticle trajectory analysis is another homogeneous method for analyzing the interaction between aptamers and viral particles. Although this method does not determine the dissociation constant, it can visualize the fraction of bound viral particles and eliminate the possibility of interaction with soluble components such as p54 protein outside viral particles. The methods requires labeling with high-yield fluorophores, like Alexa488. In this experiment, p54_5 had the highest fraction of bound virions at 20% (Table 2); we suppose that Alexa488-labeling affected the results of analysis significantly, as recombinant protein showed minor differences in this buffer.

In general, there was a good correlation found between the affinity of aptamers to recombinant p54 and ASFV virions. However, in case of Stavropol_01/08, the virus binding was disrupted due to steric reasons or variability in p54 within the virions. These events may be associated with alterations in amino acid sequence or glycosylation pattern, as it was shown for other viruses [35].

### DNA APTAMERS HAVE ANTIVIRAL ACTIVITY

Next, we tested whether the aptamers were capable of inhibiting viral infection. Vero cell line was infected with BA71V strain of ASFV using two protocols: i) co-treatment with the aptamer and ASFV, and ii) ASFV infection followed by aptamer treatment 16 hours later. The first protocol assessed the possibility of direct inhibition of viral entry into the cell. In this case, only p54_1 showed antiviral activity, with a 2-fold decrease in viral titer (Figure 3A). The second protocol assessed the assembly of new viral particles inside the cells. In this case, all aptamers showed antiviral activity, with 2-fold (p54_3) or 3-fold (p54_1 and p54_5) decreases in viral titer (Figure 3B). p54_8, with low affinity to p54, had no antiviral activity. Concluding this part, DNA aptamers targeting p54 can be used as antiviral agents.

### CONFORMATIONAL SWITCHES AND UNUSUAL NI^2+^-BINDING MOTIFS OF DNA APTAMERS

The secondary structures of the three aptamers were predicted using the RNAfold server [36] (Supplementary figure 6). The p54_1 aptamer was found to be the most structured, with its internal hairpin not assembled in the structure with minimal energy due to the formation of two alternative hairpins. This part of the structure may rearrange under certain conditions. The p54_3 aptamer contained no more than 5 pairs of nucleotides with low structuring and a shortened internal hairpin. The p54_5 aptamer had an internal hairpin, but the randomized loops are unstructured and enriched with adenines.

The aptamers were investigated using spectroscopic techniques. Circular dichroism (CD) spectroscopy is a commonly used method for studying non-canonical structures. According to the CD data, p54_1, p54_3, and p54_5 did not contain G-quadruplexes, i-motifs, or extended B-form or A-form double helices (Figure 4). A high-amplitude peak at 220 nm in the spectra of p54_3 and p54_5 corresponded to polyA inserts [37]. The consensus sequence exhibited the highest intensity for this band (Supplementary figure 7), whereas the p54_1 spectrum did not show any specific features. In the most cases CD spectra and thermal stabilities of the structures depended on the modifications introduced (Figure 4D, Table 4, Supplementary figure 8), highlighting the impact of modifications on the structural features of the aptamer. We tested several combinations of aptamer-modification-buffer that had the highest affinity to the virus in different assays. Overall, biotin destabilized the structures, Alexa over stabilized them, whereas FAM caused minor changes. The later one is preferable for label-dependent studies.

**Table 4.**
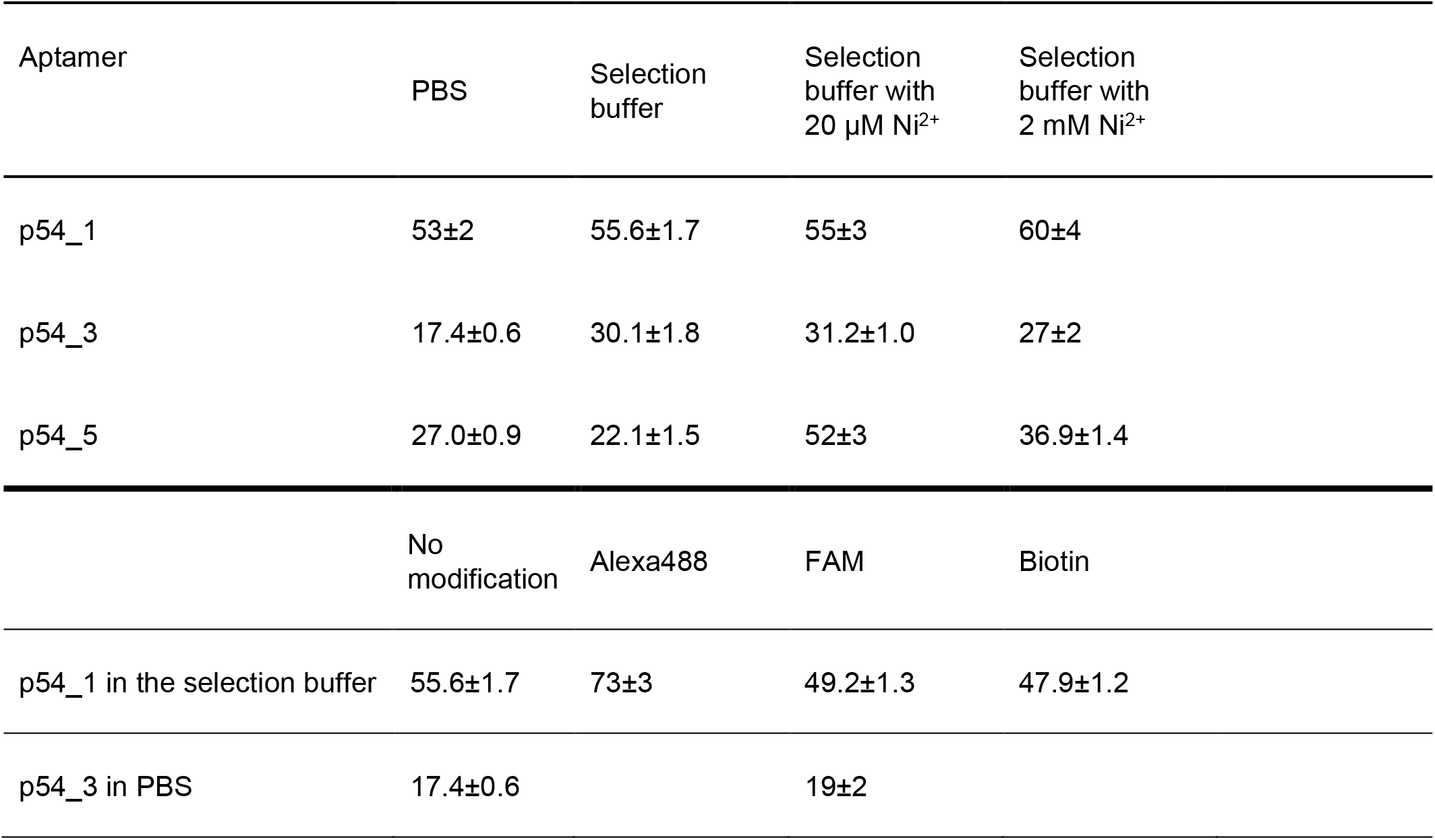
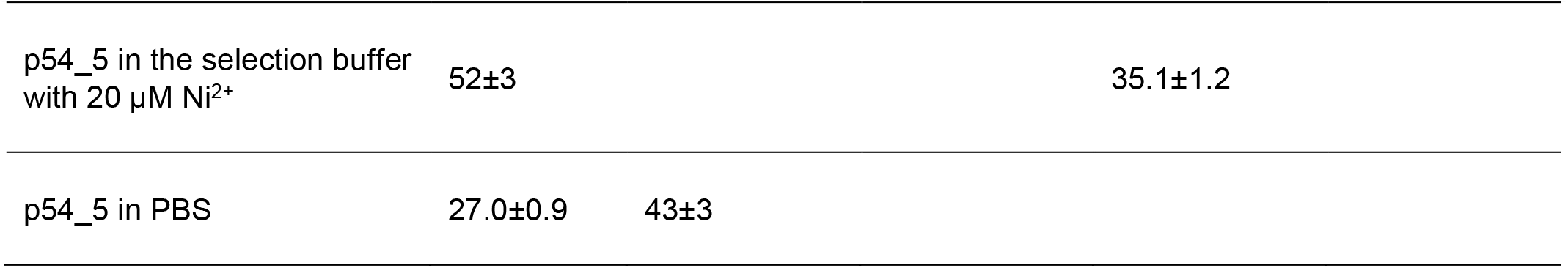
Melting temperatures of the aptamers for p54 as determined by circular dichroism spectroscopy. The values are shown in °C.

The addition of Ni^2+^ did not have a significant effect on the type of CD spectra, but the stability of p54_5 increased dramatically, with a melting temperature increase from 22°C to 52°C (Table 4, Supplementary figures 9 and 10). Stopped-flow kinetic experiments were used to determine the stoichiometry and *K*_*D*_ of aptamer-Ni^2+^ complexes. FAM-labeled aptamers were employed in this study; the changes in aptamer conformation affect the environment of the FAM-label and, hence, its fluorescence intensity [38]. p54_3 did not bind to Ni^2+^, showing no concentration dependence of the curves, while p54_1 and p54_5 readily bound Ni^2+^ (Supplementary figure 11). p54_1 had two unequal binding sites for Ni^2+^, with the first *K*_*D*_ of 0.71±0.15 µM, and the second *K*_*D*_ of 400±80 µM (Supplementary figure 12). These data explained the step-by-step increase in p54_1 affinity when increasing the concentration of Ni^2+^ from 20 µM to 2 mM (Table 1). As for p54_5, its data were approximated using a model with three equal binding sites for both Ni^2+^ (Supplementary figure 12). The estimated *K*_*D*_ value was 13±3 µM.

We choose p54_1 for the further structural analysis aiming to identify structural base of affinity changes due to Ni^2+^ coordination. Molecular modeling (MM), small angle X-ray scattering (SAXS) and ^1^H NMR spectroscopy were employed to uncover conformational switching in the aptamer. The AlphaFold 3 program was used to create the 3D model; its capabilities for predicting the 3D structure of aptamers, including non-canonical forms, are presented in the article [39]. In addition to the models obtained using AlphaFold 3, models were built using the strategy developed by Tsvetkov [40] for non-canonical forms adapted for aptamers. Next, each model was tested for stability using the molecular dynamics method. Based on the calculated data, the energy contributions, the number of hydrogen bonds, the energy of interaction of aptamers with ions, and the stability of ion localization were estimated (Supplementary figures 13, 14). Six stable models with comparable energies and number of H-bonds were created for p54_1 without Ni^2+^, also, seven stable models were created for p54_1 with Ni^2+^. All models have duplex regions formed by complementary pairs. For p54_1 without cations, one of the possible structures contains a hybrid G-quadruplex, others are DNA hairpins with the following AT/GC pair ratios: 1^st^ - 4:10, 2^nd^ - 4:9, 3^rd^ - 4:10, 4^th^ - 2:4, 5^th^ - 3:5, 6^th^ - 2:4.

^1^H NMR spectroscopy was used in an attempt to select the correct model (Supplementary Figure 15). The spectra showed that the p54_1 aptamer had weak G-quadruplex-related signals, but their intensities were too low to support an assembled G-quadruplex structure. Watson–Crick base pairs were observed, with an AT/GC pair ratio close to 3:5. The fifth model, with an AT/GC pair ratio of approximately 3:5, is the closest to this observation. In addition, the fifth model contains several G–G interactions that could account for the low-intensity signals observed in the G-quadruplex region. The addition of 2 equivalents of Ni^2+^ did not noticeably change the ^1^H NMR spectra. The only observable change was signal broadening, which increased upon addition of Ni ions. Zn^2+^ ions did not cause broadening and therefore produced no observable changes in the spectra.

SAXS was used as additional technique that can identify high-amplitude conformational changes. SAXS data definitively confirmed the cation-induced conformational transition and structural compaction of the p54_1. Upon the addition of Ni^2+^ ions, a distinct increase in the maximum particle diameter (D_max_) from 55.0 to 65.0 Å and the radius of gyration (R_g_) from 19.66 to 20.57 Å was observed (Supplementary Table 3). This behavior was further substantiated by the visual analysis of the pair-distance distribution function P(r). For the cation-free state, the P(r) profile exhibits a prominent asymmetric peak skewed toward smaller distances with a relatively steep decay, which is characteristic of an L-shaped, bent ‘boomerang’ conformation where the mass is concentrated around the central hinge point (Figure 5, Supplementary Figure 16). Conversely, upon nickel binding, the P(r) curve broadens significantly and transitions into a classic elongated profile with a smooth, extended tail that gradually decays to zero at 65.0 Å. This profile shape strictly indicates a cation-induced straightening of the aptamer backbone, transitioning from a highly bent conformation into an elongated, rod-shaped structure (Figure 5, Supplementary Figure 17). Notably, despite the increase in longitudinal length, the Porod volume (V_p_) decreases substantially by ∼1512 Å^3^ upon nickel binding (from 20420 to 18908 Å^3^). This phenomenon provides evidence on more dense packing of the structure in the Ni^2+^-complex.

Based on the data from ^1^H NMR and SAXS, the most appropriate MM models were chosen. 5^th^ model is very similar to the aptamer without Ni^2+^, while 7^th^ model with Ni^2+^ corresponds to the aptamer with 2 Ni^2+^ cations. Detailed comparison of polar contact sets of p54_1 without of Ni^2+^ and with Ni^2+^ is shown in Supplementary table 4. The conformational changes during the transition between two models are significant. The loops changed their position, as illustrated in Figure 5. Hairpin stems were preserved, while loop packing was changed completely. Both Ni^2+^ ions are coordinated close to each other. Each Ni^2+^ ion is coordinated with one phosphate group and oxygen of cytosine that caused distortion of the aptamer backbone rearranging loops into a new form (Figure 5).

This Ni^2+^-dependent interface is rather unusual. Previously, structures of individual DNA-Ni^2+^ complexes were not described. However, Ni^2+^-dependent DNAzymes were described [41,42]. The activity of these DNAzymes is turned on in the presence of Ni^2+^ and turned off without these cations. The oligonucleotides were selected from a library made up of duplex with a large randomized loop. This loop have no direct sequence similarity with p54_1, however, the overall structuring might have some similarity. In this case, Ni^2+^ can be used as a tiny regulator of DNA structure and function in artificial constructions that can be used, for example, as switchable recognition elements for affinity chromatography or biosensors.

## Conclusion

This study reports for the first time on DNA aptamers that target the p54 viral protein of ASFV. The aptamers bound to both recombinant protein and ASFV particles as well as exhibited antiviral properties. They can be used as both recognition elements and inhibitory compound. Additionally to this basic results, we identified the aptamer whose affinity depended on Ni^2+^. Molecular modeling, ^1^H NMR and SAXS were used to identify the Ni^2+^-dependent conformational switch of this aptamer. p54_1 has several possible conformations with close energies, so it is an example of conformational heterogeneous structure. Ni^2+^ caused bending of the phosphate backbone stabilizing one of the possible conformation of the loops that significantly affects affinity. Cation-dependent regulation of aptamer activity is promising as it allows regeneration of affine carriers and biosensors with chelating agents, while addition of Ni^2+^ restores the high-affine aptamer conformation.

## Supporting Information

The authors have cited additional references within the Supporting Information.^[43-46]^

## Acknowledgements

This work was supported by JINR project 07-5-1131-3-2025/2029. NMR experiments were performed with financial support from the State Assignment #124012400364-9 of Lomonosov Moscow State University. The stopped flow experiments were performed using the equipment of the MSU Shared Research Equipment Center “Subdiffractional microscopy and spectroscopy” and purchased by the MSU as part of the Equipment Renovation Program (National Project “Science”). The circular dichroism experiments were performed using the equipment of the MSU Shared Research Equipment Center “Subdiffraction microscopy and spectroscopy” and purchased by MSU in the frame of the Equip-ment Renovation Program (National Project “Science”).

## Entry for the Table of Contents

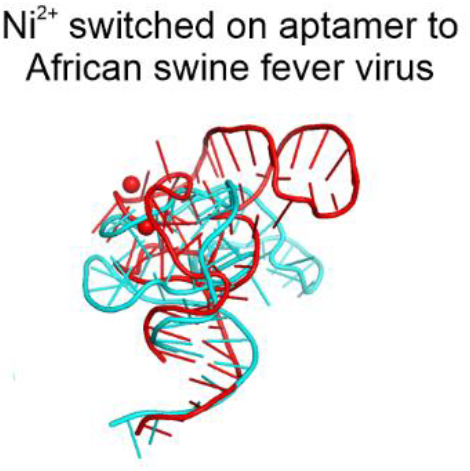

## Notes

Supporting information for this article is given via a link at the end of the document.

### Competing Interest Statement

The authors have declared no competing interest.

### Summary of Updates

Structural study of p54_1 aptamer was added

