## Supplementary for "ARTIFICIAL RECOGNITION ELEMENTS FOR AFRICAN SWINE FEVER VIRUS: DNA APTAMERS WITH UNUSUAL NI^2+^-BINDING MOTIFS"

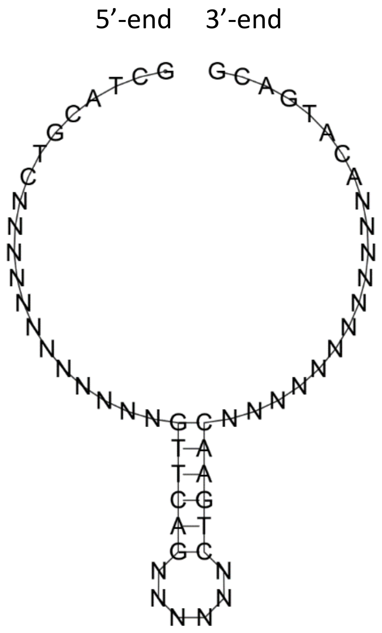


**Supplementary figure 1.** The structure of the randomized library with an inserted DNA hairpin.

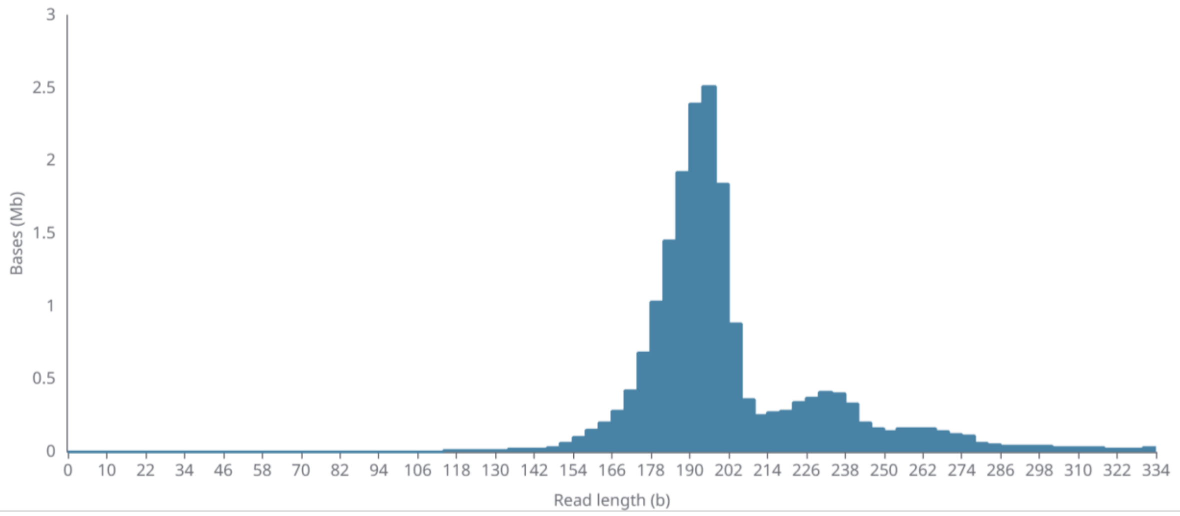


**Supplementary figure 2.** Analysis of the length of sequenced fragments estimated using MinKNOW software.

**Supplementary table 1.** The alignment results for 3,380 sequences obtained from a single selection cycle, with the numbers reflecting the percentage probability of nucleotide appearance.

| Position | A | T | G | C | Deletion | Consensus |
|---|---|---|---|---|---|---|
| 1 | 0 | 0 | 100 | 0 | 0 | G |
| 2 | 0 | 0 | 0 | 100 | 0 | C |
| 3 | 0 | 100 | 0 | 0 | 0 | T |
| 4 | 100 | 0 | 0 | 0 | 0 | A |
| 5 | 0 | 0 | 0 | 100 | 0 | C |
| 6 | 0 | 0 | 100 | 0 | 0 | G |
| 7 | 0 | 100 | 0 | 0 | 0 | T |
| 8 | 0 | 0 | 0 | 100 | 0 | C |
| 9 | 69 | 3 | 8 | 20 | 1 | A |
| 10 | 72 | 5 | 7 | 15 | 1 | A |
| 11 | 67 | 4 | 9 | 19 | 1 | A |
| 12 | 68 | 4 | 7 | 16 | 5 | A |
| 13 | 67 | 4 | 8 | 17 | 4 | A |
| 14 | 67 | 4 | 8 | 17 | 4 | A |
| 15 | 64 | 4 | 8 | 18 | 6 | A |
| 16 | 56 | 5 | 8 | 19 | 13 | A |
| 17 | 62 | 4 | 8 | 15 | 10 | A |
| 18 | 61 | 4 | 7 | 15 | 13 | A |
| 19 | 59 | 4 | 6 | 15 | 17 | A |
| 20 | 61 | 3 | 8 | 11 | 17 | A |
| 21 | 6 | 3 | 81 | 2 | 8 | G |
| 22 | 2 | 84 | 3 | 2 | 9 | T |
| 23 | 1 | 86 | 1 | 4 | 8 | T |
| 24 | 3 | 5 | 1 | 83 | 8 | C |
| 25 | 87 | 1 | 3 | 2 | 7 | A |
| 26 | 6 | 1 | 84 | 2 | 8 | G |
| 27 | 59 | 6 | 9 | 12 | 13 | A |
| 28 | 61 | 6 | 8 | 14 | 12 | A |
| 29 | 60 | 5 | 10 | 15 | 10 | A |
| 30 | 63 | 6 | 9 | 14 | 10 | A |
| 31 | 64 | 6 | 8 | 14 | 8 | A |
| 32 | 68 | 6 | 6 | 11 | 9 | A |
| 33 | 2 | 3 | 1 | 86 | 8 | C |
| 34 | 1 | 87 | 2 | 2 | 8 | T |
| 35 | 4 | 2 | 85 | 1 | 8 | G |
| 36 | 86 | 1 | 2 | 1 | 11 | A |
| 37 | 88 | 1 | 2 | 2 | 7 | A |
| 38 | 3 | 1 | 1 | 85 | 10 | C |
| 39 | 60 | 6 | 9 | 11 | 14 | A |
| 40 | 57 | 8 | 11 | 12 | 12 | A |
| 41 | 54 | 7 | 9 | 10 | 20 | A |
| 42 | 57 | 8 | 11 | 11 | 13 | A |
| 43 | 57 | 8 | 11 | 11 | 12 | A |
| 44 | 58 | 8 | 11 | 12 | 12 | A |
| 45 | 58 | 9 | 12 | 12 | 10 | A |
| 46 | 57 | 10 | 12 | 13 | 8 | A |
| 47 | 54 | 11 | 15 | 12 | 8 | A |
| 48 | 48 | 13 | 18 | 14 | 8 | A |
| 49 | 47 | 14 | 18 | 14 | 8 | A |
| 50 | 45 | 14 | 17 | 17 | 7 | A |
| 51 | 94 | 1 | 2 | 1 | 3 | A |
| 52 | 1 | 1 | 1 | 93 | 4 | C |
| 53 | 92 | 1 | 2 | 1 | 5 | A |
| 54 | 1 | 94 | 1 | 1 | 2 | T |
| 55 | 3 | 2 | 91 | 1 | 3 | G |
| 56 | 92 | 3 | 1 | 1 | 2 | A |
| 57 | 1 | 4 | 1 | 92 | 2 | C |
| 58 | 4 | 2 | 90 | 1 | 4 | G |

**Supplementary table 2.** Sequences of DNA oligonucleotides studied in the work. N is a random nucleotide, non–randomized nucleotides are underlined. AF488 is an Alexa dye with an excitation wavelength of 488 nm, and FAM is carboxyfluorescein.

| Code | Sequences (5´→3´) |
|---|---|
| Initial library | GCTACGTC NNNNNNNNNNNN GTTCAG NNNNNN CTGAAC NNNNNNNNNNNN ACATGACG |
| Consensus | GCTACGTC AAAAAAAAAAAA GTTCAG AAAAAA CTGAAC AAAAAAAAAAAA ACATGACG |
| p54_1 | GCTACGTC GGGACAGGGGAA TTCAG GAGTCA CTGAAC AGCAGGCGCC ACATGACG |
| p54_2 | GCTACGTC AAAAAACACAA GTTCAG AACACC CTGAAC AAGGGAAAGAAC ACATGACG |
| p54_3 | GCTACGTC GAATACAAAACA G AG AAGGAA CTCAAC AAAAAAAGAAAC ACATGACG |
| p54_4 | GCTACGTC CAAAATA GTTCAG AAAAAA CTGAAC AAAAACAAAAAG ACATGACG |
| p54_5 | GCTACGTC AACAAAAAAACA GTTCAG AAAACA CTGAAC CAGGAAAAGAAC ACATGACG |
| p54_6 | GCTACGTC GACGGGC GTTCAG CACACA CTGAAC AAGGCGGCGGGG ACATGACG |
| p54_7 | GCTACGTC CAGGGGAA GTT GAAGTG C ACATGACG |
| p54_8 | GCTACGTC AACAAAAATAAT GTTCAG TAGACA CTGAAC AGCCATGC ACATCAAT |
| p54_9 | GCTACGTC CACCCCAAAATA GTTCAG TATTAG CTGAAC CTACAAACGCC ACATGACG |
| p54_10 | GCTACGTC CAACAAAAAGAG GTTCAG AAATAA CTGAAC GAACCCCCCCA ACATGACG |
| p54_1_AF | GCTACGTC GGGACAGGGGAA TTCAG GAGTCA CTGAAC AGCAGGCGCC ACATGACG [AF488] |
| p54_3_AF | GCTACGTC GAATACAAAACA G AG AAGGAA CTCAAC AAAAAAAGAAAC ACATGACG [AF488] |
| p54_5_AF | GCTACGTC AACAAAAAAACA GTTCAG AAAACA CTGAAC CAGGAAAAGAAC ACATGACG [AF488] |
| p54_8_AF | GCTACGTC AACAAAAATAAT GTTCAG TAGACA CTGAAC AGCCATGC ACATCAAT [AF488] |
| p54_1_Bio | [Biotin] GCTACGTC GGGACAGGGGAA TTCAG GAGTCA CTGAAC AGCAGGCGCC ACATGACG |
| p54_3_Bio | [Biotin] GCTACGTC GAATACAAAACA G AG AAGGAA CTCAAC AAAAAAAGAAAC ACATGACG |
| p54_5_Bio | [Biotin] GCTACGTC AACAAAAAAACA GTTCAG AAAACA CTGAAC CAGGAAAAGAAC ACATGACG |
| p54_8_Bio | [Biotin] GCTACGTC AACAAAAATAAT GTTCAG TAGACA CTGAAC AGCCATGC ACATCAAT |
| p54_1_FAM | GCTACGTC GGGACAGGGGAA TTCAG GAGTCA CTGAAC AGCAGGCGCC ACATGACG [FAM] |
| p54_3_FAM | GCTACGTC GAATACAAAACA G AG AAGGAA CTCAAC AAAAAAAGAAAC ACATGACG [FAM] |
| p54_5_FAM | GCTACGTC AACAAAAAAACA GTTCAG AAAACA CTGAAC CAGGAAAAGAAC ACATGACG [FAM] |
| p54_8_FAM | GCTACGTC AACAAAAATAAT GTTCAG TAGACA CTGAAC AGCCATGC ACATCAAT [FAM] |

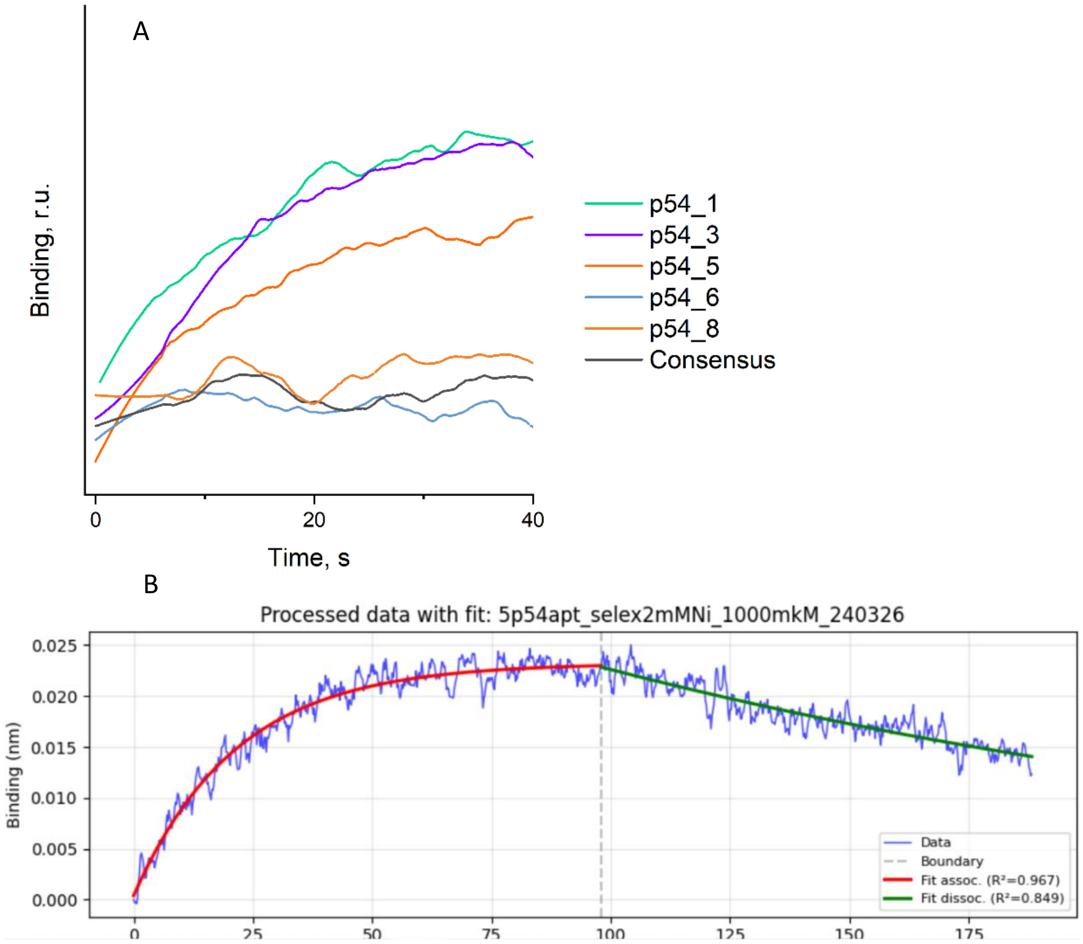


**Supplementary figure 3.** Affinity estimation of aptamers for p54 protein using biolayer interferometry in the selection buffer (a) and an example of curve fitting using the BliMart software for aptamer p54_5 in the selection buffer with 20 µM $NiCl_2$.

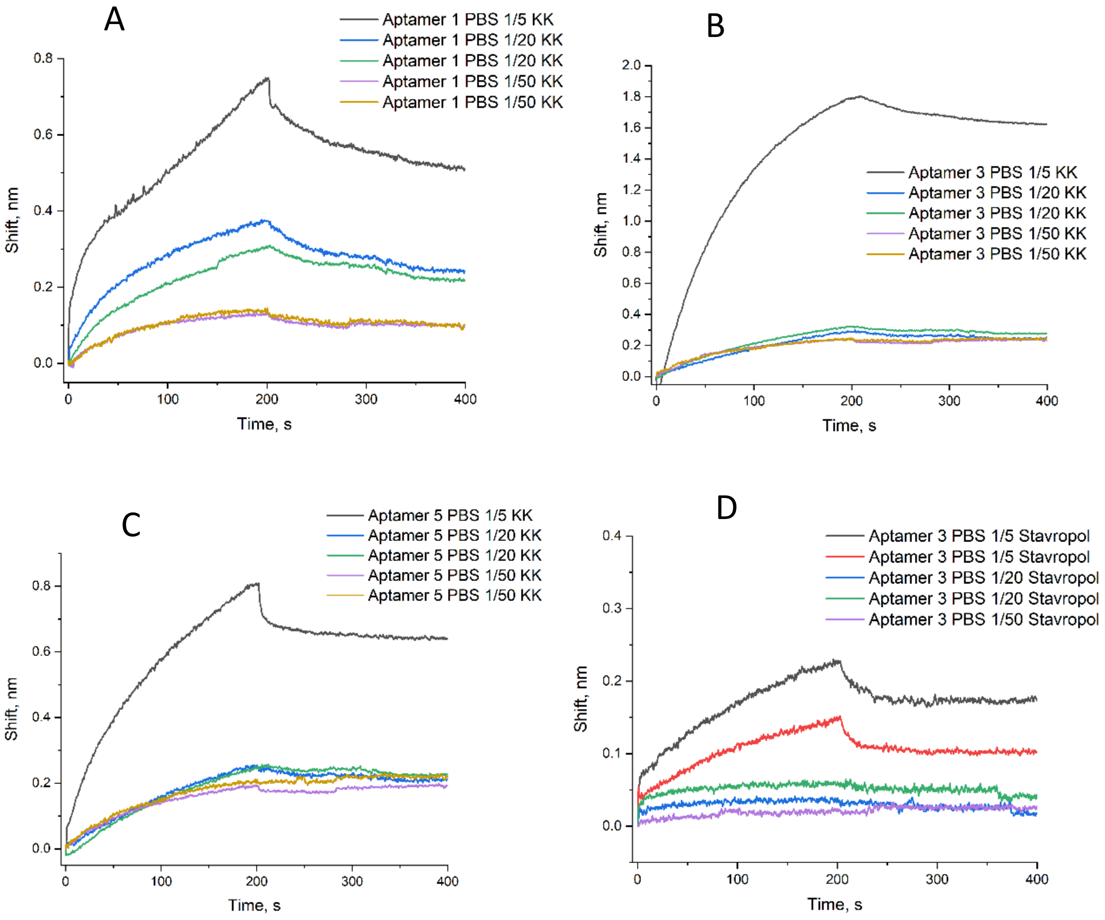


**Supplementary figure 4.** Affinity estimation of ASFV virions using biolayer interferometry. The curves for biotinylated aptamers p54_1(A), p54_3 (B,D), and p54_5 (C) with ASFV KK262 cos-1 (A-C) and Stavropol 01/08 MF (D) in PBS.

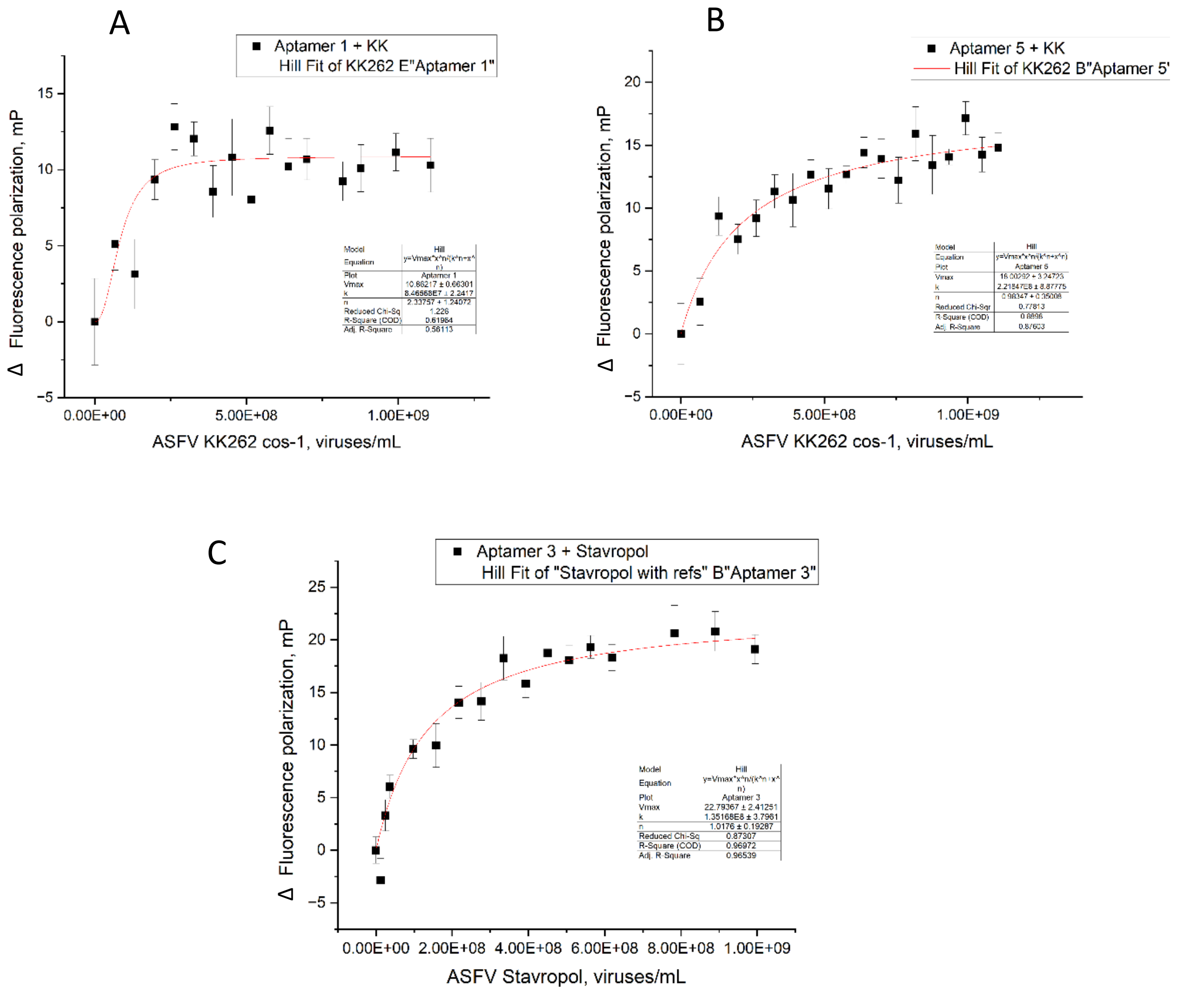


**Supplementary figure 5.** Affinity estimation of ASFV virions using fluorescence polarization assay, The dependencies for FAM-modified aptamers p54_1 (A), p54_5 (B), and p54_3 (C) with ASFV KK262 cos-1 (A,B) and Stavropol 01/08 MF (C) viruses in PBS.

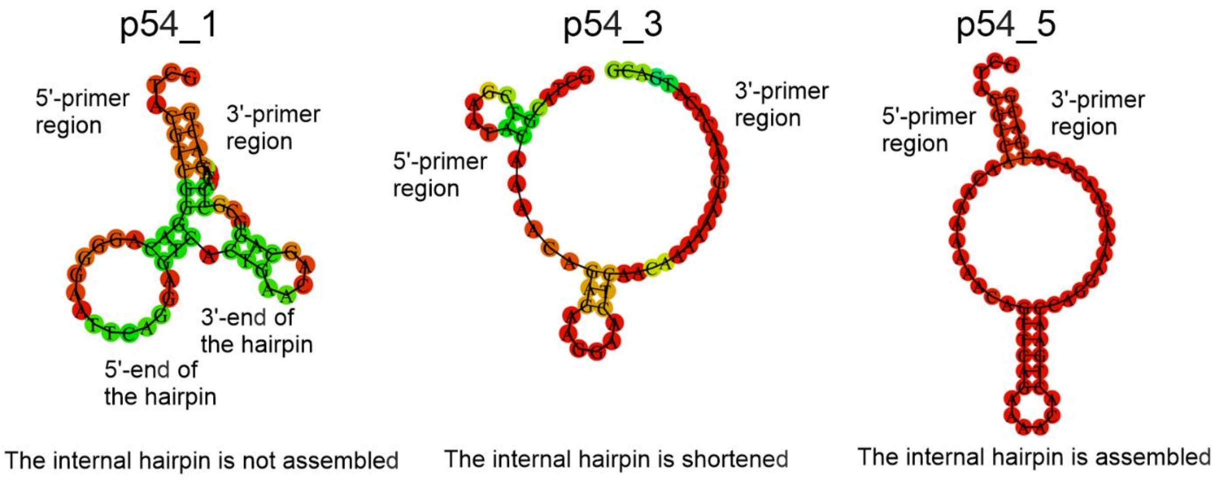


**Supplementary figure 6.** The secondary structures of the aptamers predicted by the RNAfold server.

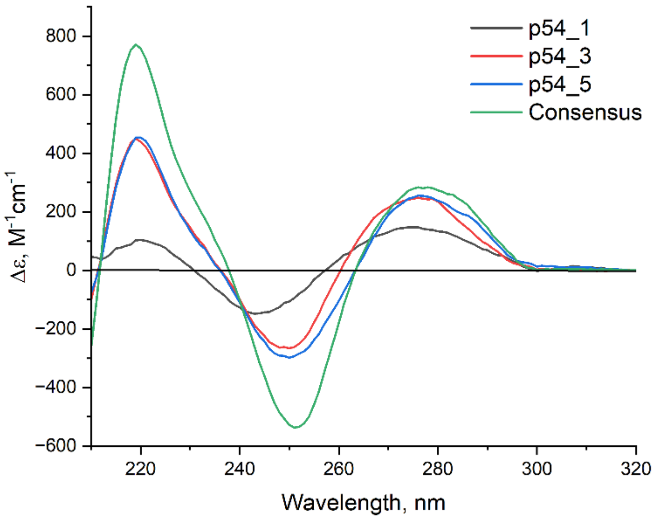


**Supplementary figure 7.** The circular dichroism spectra of the aptamers in the selection buffer containing 20 µM $NiCl_2$. The spectra were acquired for a 1 µM solution of each aptamer.

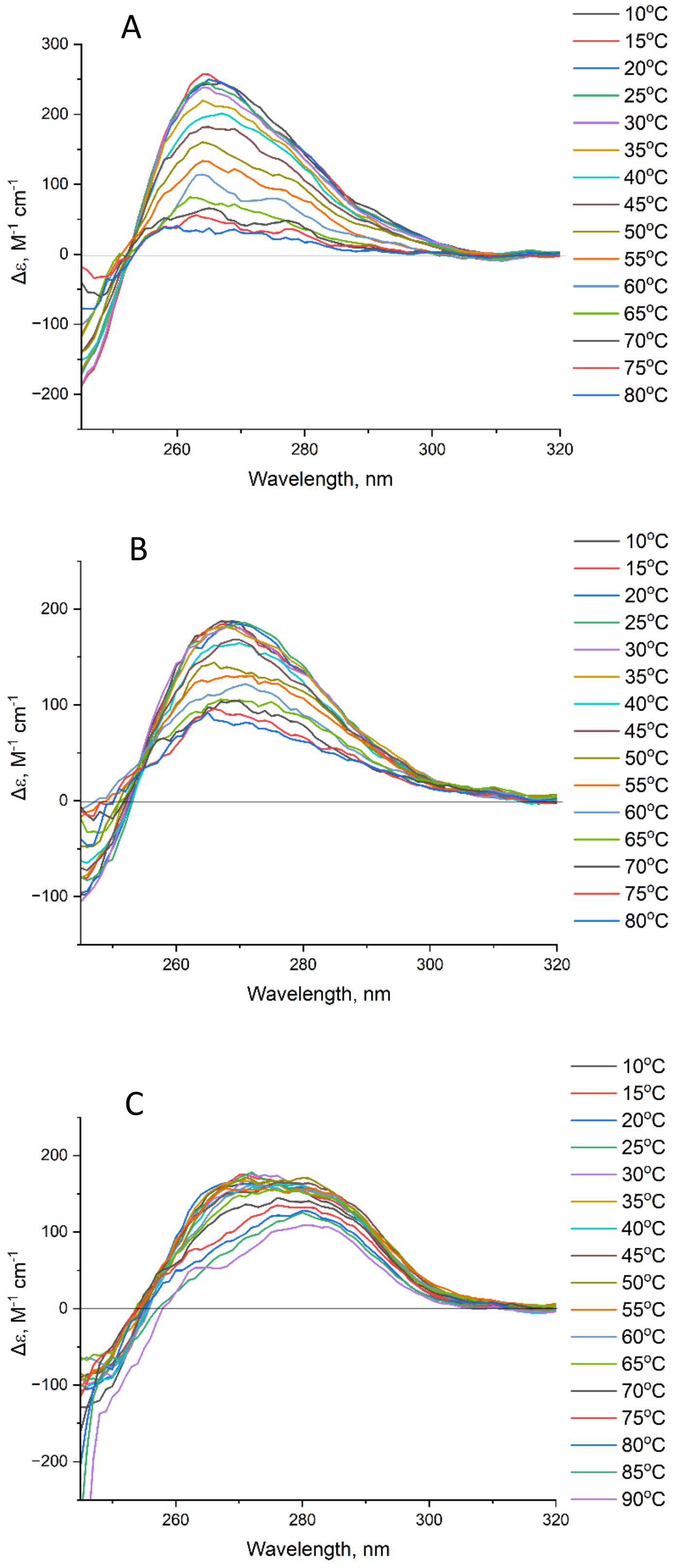


**Supplementary figure 8.** The melting experiment of circular dichroism spectroscopy for modified variants of p54_1 in the selection buffer containing 20 µM $NiCl_2$. (A) Biotin, (B) FAM, and (C) Alexa488.

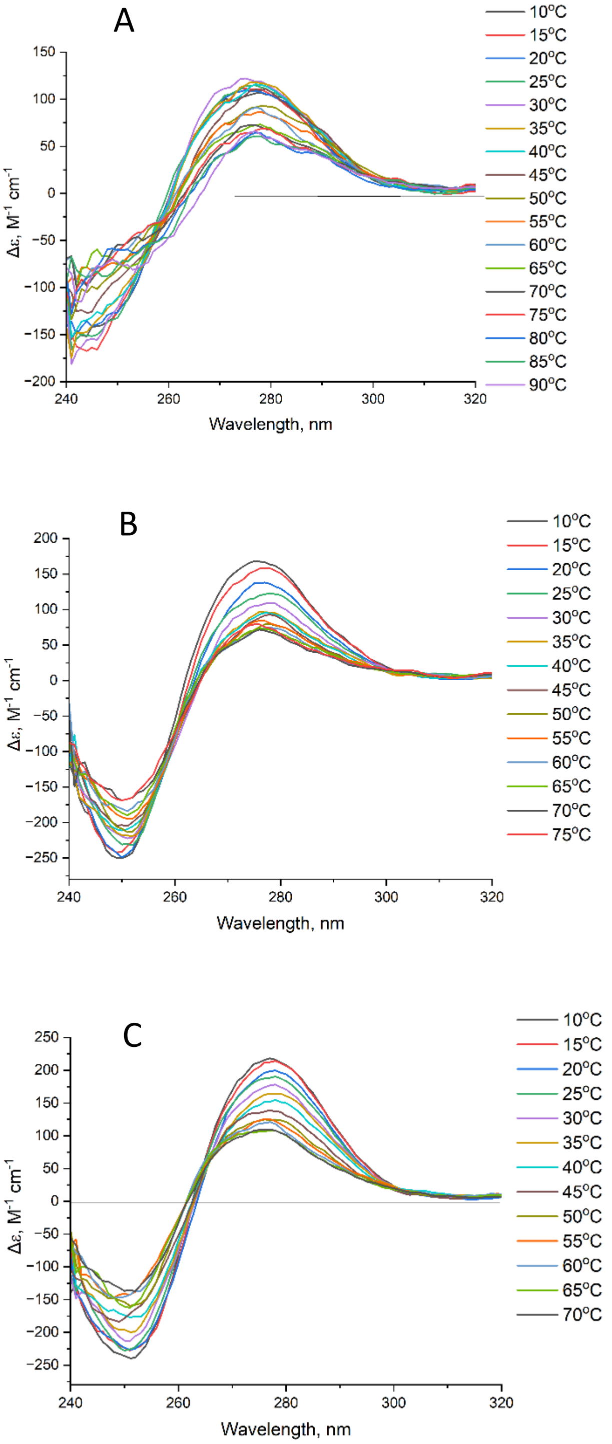


**Supplementary figure 9.** The melting experiment of circular dichroism spectroscopy of the aptamers in PBS: (A) p54_1, (B) p54_3, and (C) p54_5.

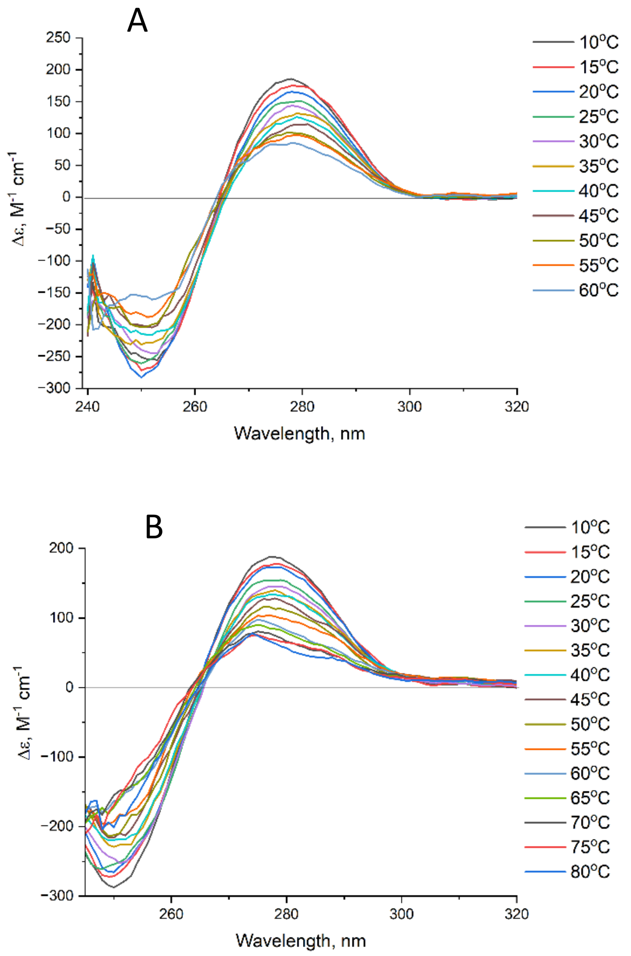


**Supplementary figure 10.** The melting experiment of circular dichroism spectroscopy of p54_5 in the selection buffer (A), the selection buffer with 20 µM $NiCl_2$ (B).

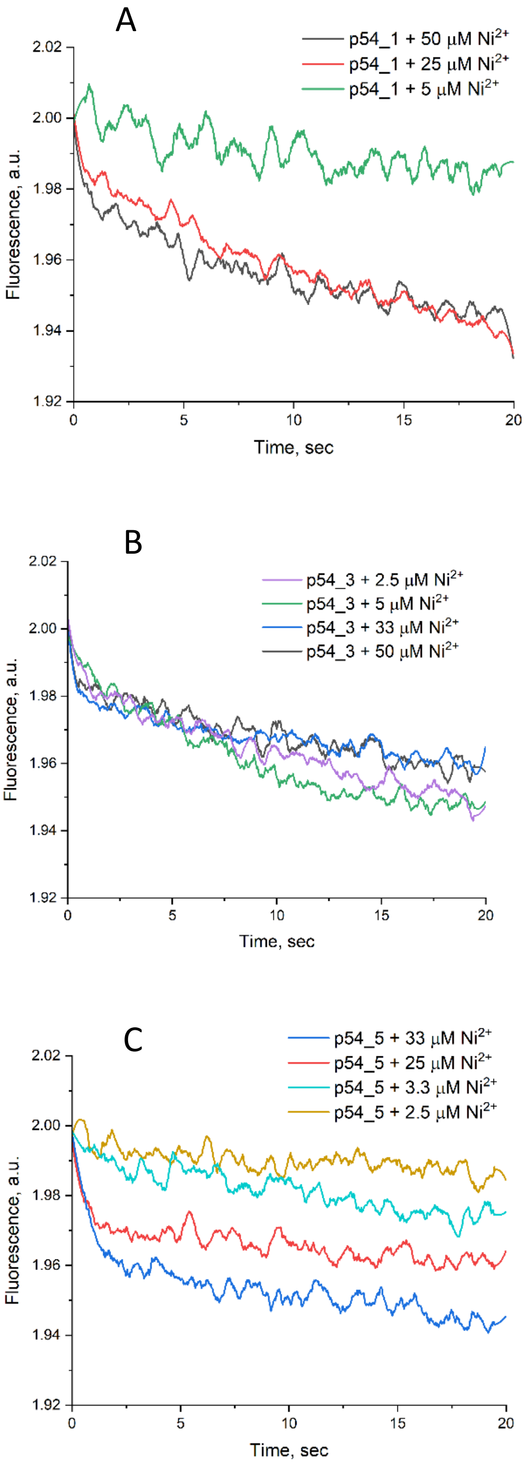


**Supplementary figure 11.** $Ni^{2+}$-dependent changes in FAM-labeled aptamers acquired within stopped-flow kinetic experiments: p54_1 (A), p54_3 (B), and p54_5 (C). Fluorescence of p54_1 and p54_5 have dependence on $Ni^{2+}$ concentration, while p54_3 has no dependence with fluorophore bleaching only.

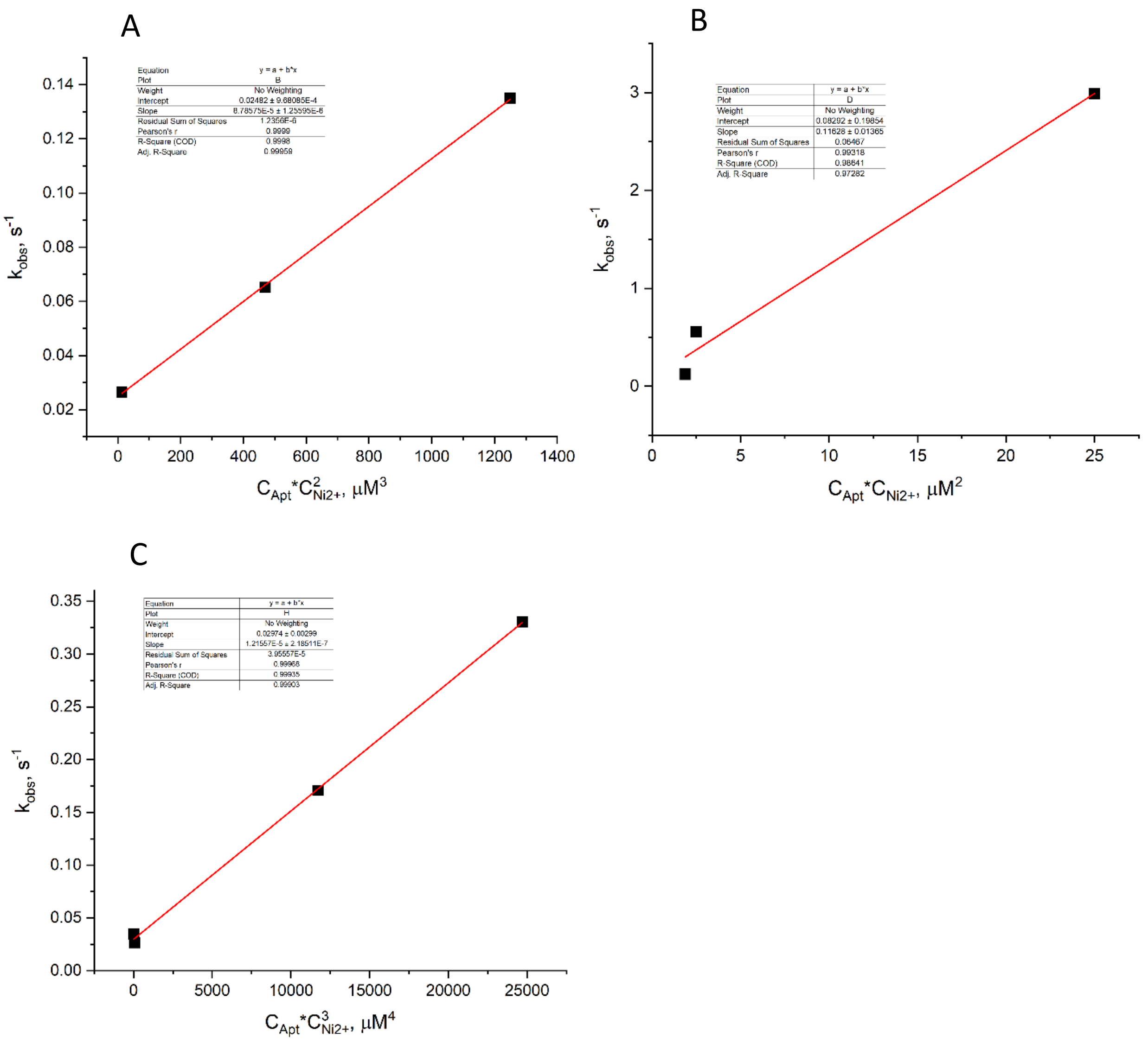


**Supplementary figure 12.** Linearization of the $k_{obs}$ values used to determine the stoichiometry and $K_D$ of aptamer-$Ni^{2+}$ complexes: (A) p54_1 with $Ni^{2+}$, a model with two binding sites; (B) p54_1 with $Ni^{2+}$, the first binding site in a model with non-equal binding sites; (C) p54_5 with $Ni^{2+}$, a model with three equal binding sites.

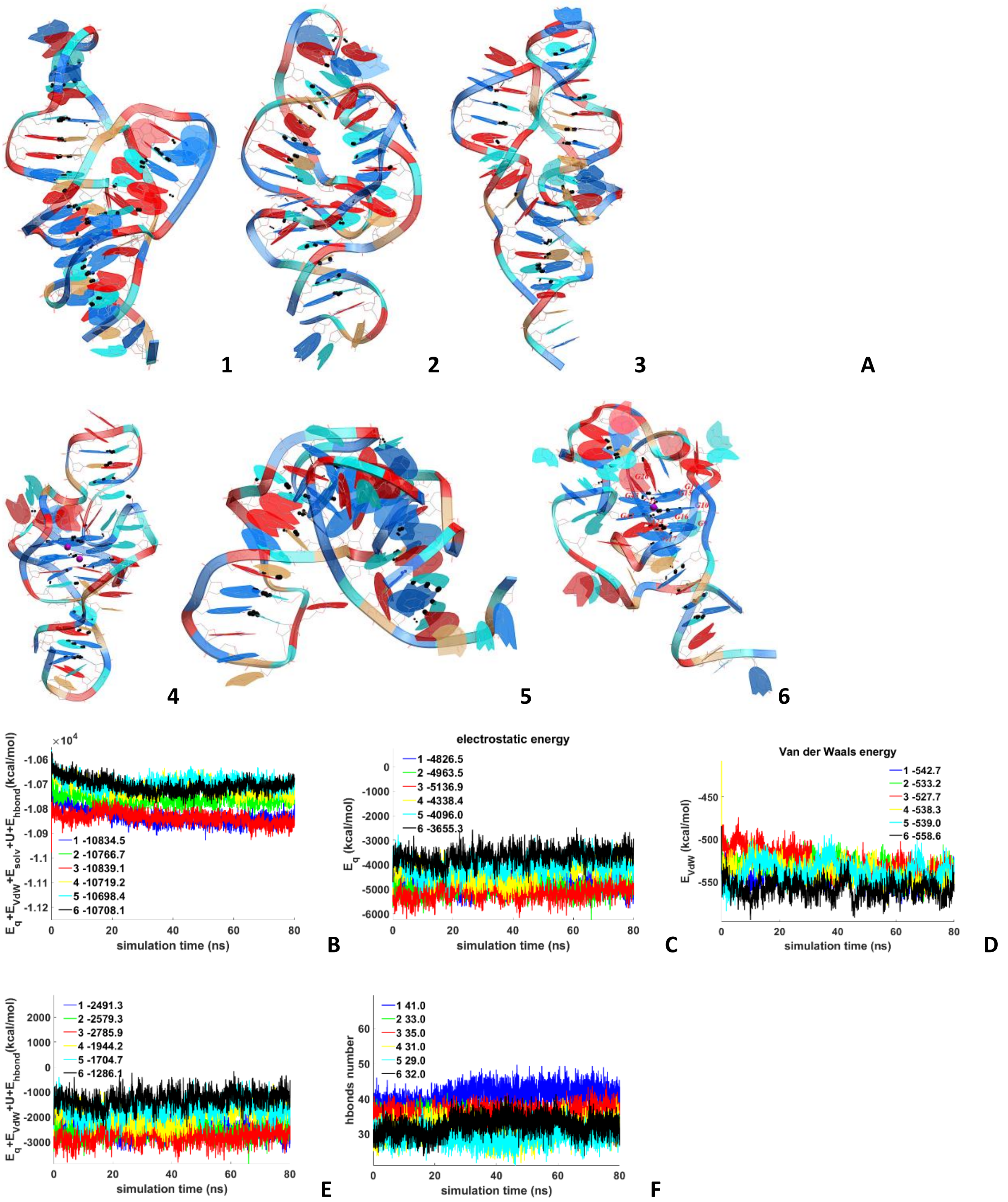


**Supplementary figure 13.** The conformation trajectories of p54_1 without $Ni^{2+}$ and the evolution of energy contributions. A – conformations of models 1–5. DNA visualization is colored as follows: G – blue, C – light blue, A – red, T – light brown. Atoms are represented by the following colors: carbon – gray; oxygen – red; nitrogen – blue; hydrogen – white (only polar hydrogens are shown); nickel ions – purple; potassium ions – magenta. Hydrogen bonds are shown as dotted lines; the thickness depends on the bond energy. B - $E_{eq}$ + $E_{vdw}$ + $E_{solv}$ + U + $E_{hbond}$; $E_{eq}$ – electrostatic, $E_{vdw}$ -Van der Waals, $E_{solv}$=$E_{GB}$ + $E_{surf}$, $E_{GB}$ - polar energy of solvation and $E_{surf}$ - non-polar energy of solvation due to the hydrophobic surface available to the solvent; U - stress energy of the bonds, valence and dihedral angles; $E_{hbond}$ - hydrogen bond energy. B – C - the evolution of the $E_{eq}$ and $E_{vdw}$. E - evolution of the internal energy values. F - evolution of the number of intramolecular hydrogen bondshe plots were smoothed using the moving average method (span = 5), for the H-bond numbers (span = 3). The average over the entire trajectory were in the legends.

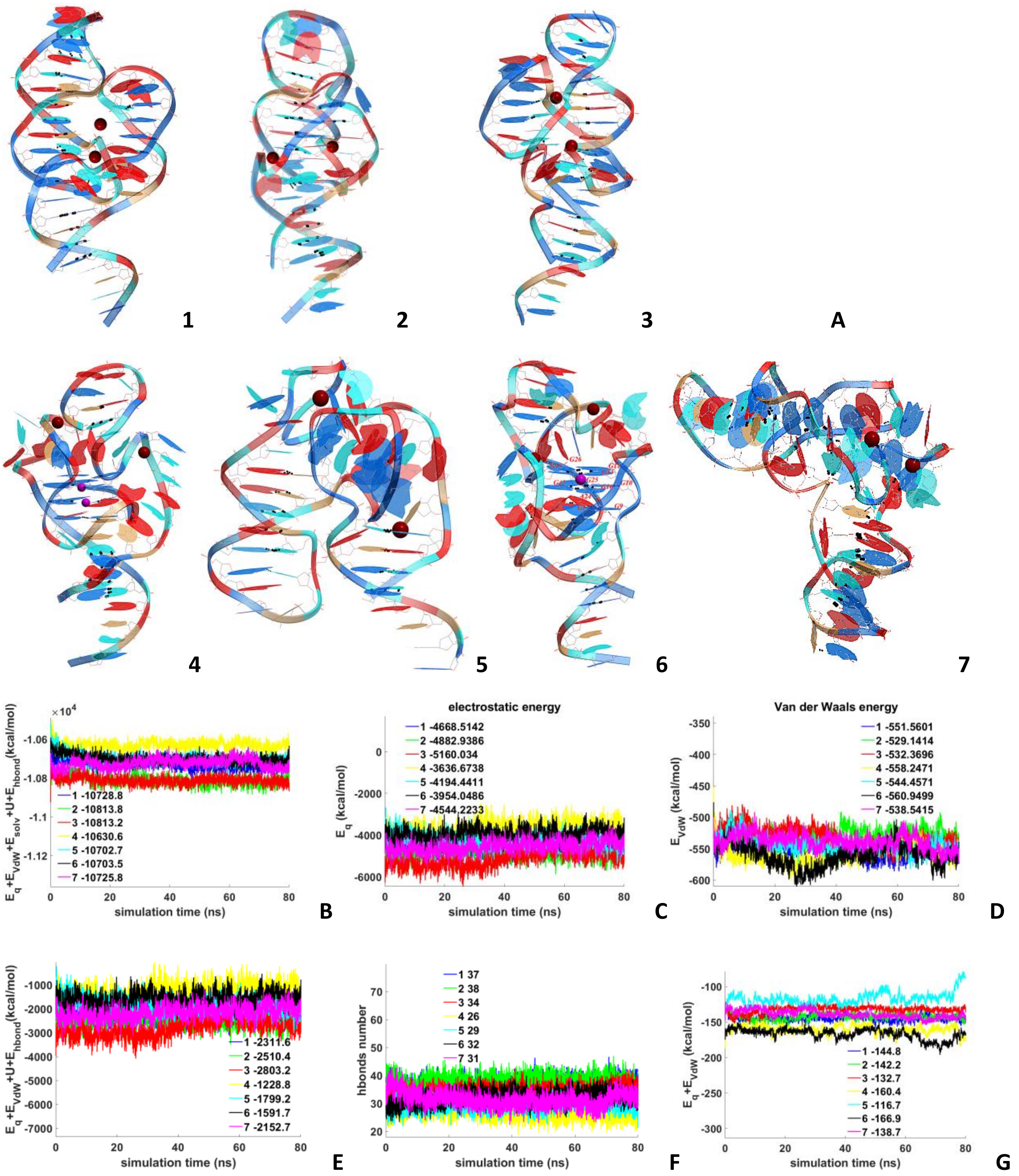


**Supplementary figure 14.** The conformation trajectories of p54_1 with 2 $Ni^{2+}$ ions and the evolution of energy contributions. A – conformations of models 1–5. DNA visualization is colored as follows: G – blue, C – light blue, A – red, T – light brown. Atoms are represented by the following colors: carbon – gray; oxygen – red; nitrogen – blue; hydrogen – white (only polar hydrogens are shown); nickel ions – purple; potassium ions – magenta. Hydrogen bonds are shown as dotted lines; the thickness depends on the bond energy. B - $E_{eq}$ + $E_{vdw}$ + $E_{solv}$ + U + $E_{hbond}$; $E_{eq}$ – electrostatic, $E_{vdw}$ -Van der Waals, $E_{solv}$=$E_{GB}$ + $E_{surf}$, $E_{GB}$ - polar energy of solvation and $E_{surf}$ - non-polar energy of solvation due to the hydrophobic surface available to the solvent; U - stress energy of the bonds, valence and dihedral angles; $E_{hbond}$ - hydrogen bond energy. B – C - the evolution of the $E_{eq}$ and $E_{vdw}$. E - evolution of the internal energy values. F - evolution of the number of intramolecular hydrogen bonds. G - evolution of the values of the interaction energy between DNA and $Ni^{2+}$ ($E_{eq}$ + $E_{vdw}$). The plots were smoothed using the moving average method (span = 5), for the H-bond numbers (span = 3). The average over the entire trajectory were in the legends.

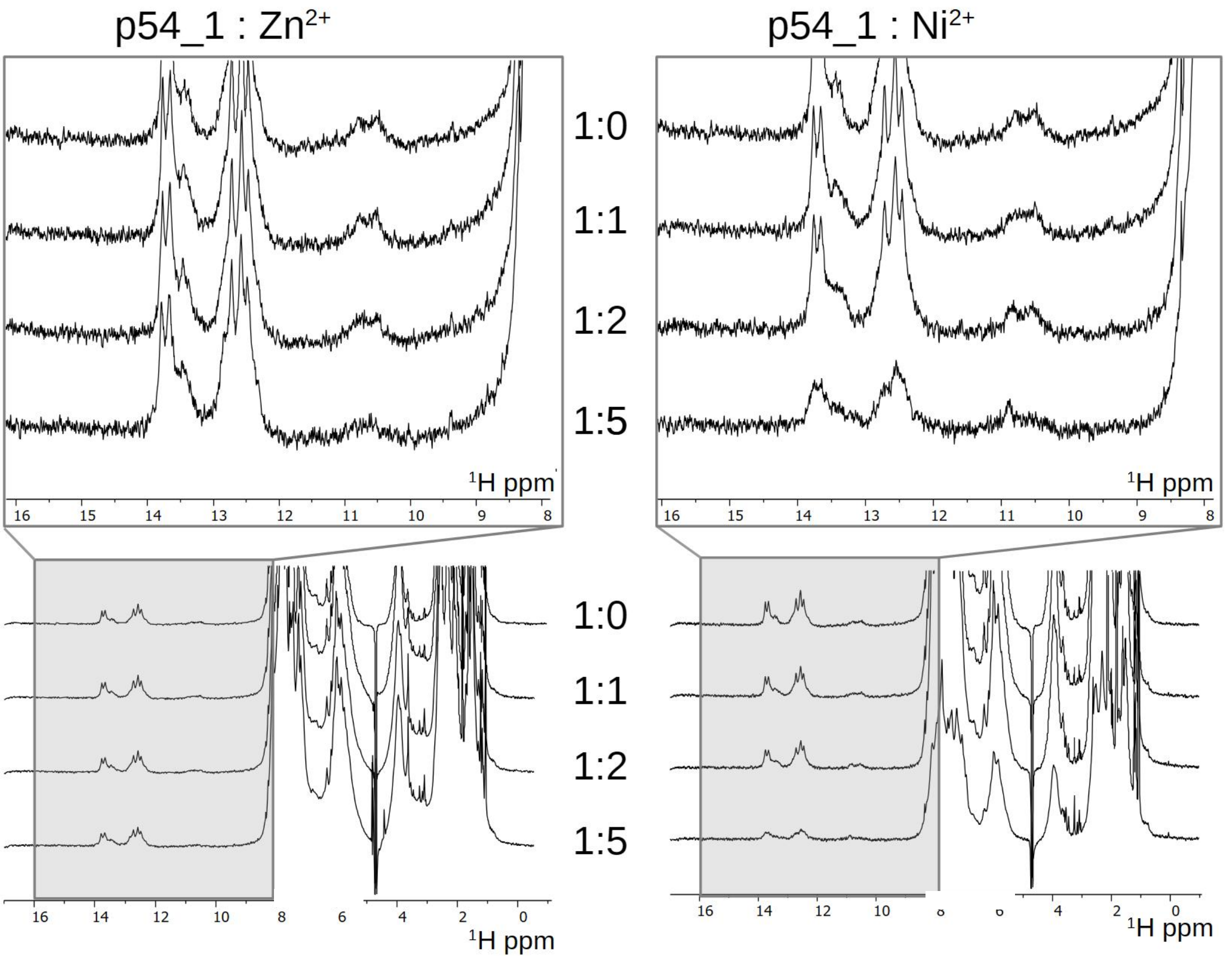


**Supplementary figure 15.** $^1H$ NMR spectra of p54_1 in the presence of increasing amounts of the target cation ($Ni^{2+}$, right panel) and the off-target cation ($Zn^{2+}$, left panel). The molar ratio between p54_1 and metal ion is indicated. The top panels show expansions of the imino proton region from the full spectra displayed in the bottom panels.

**Supplementary Table 3.** Structural parameters of the p54_1 aptamer and its complex with $Ni^{2+}$ derived from SAXS data.

| Aptamer state | $Rg$, (Å) | $Dmax$, (Å) | $Vp$, (Å$^3$) | Conformation |
|---|---|---|---|---|
| Cation-free | 19.66 | 55.0 | 20 420 | L-shaped (bent 'boomerang' conformation) |
| $Ni^{2+}$-bound | 20.57 | 65.0 | 18 908 | Extended cylinder (rod-shaped conformation) |

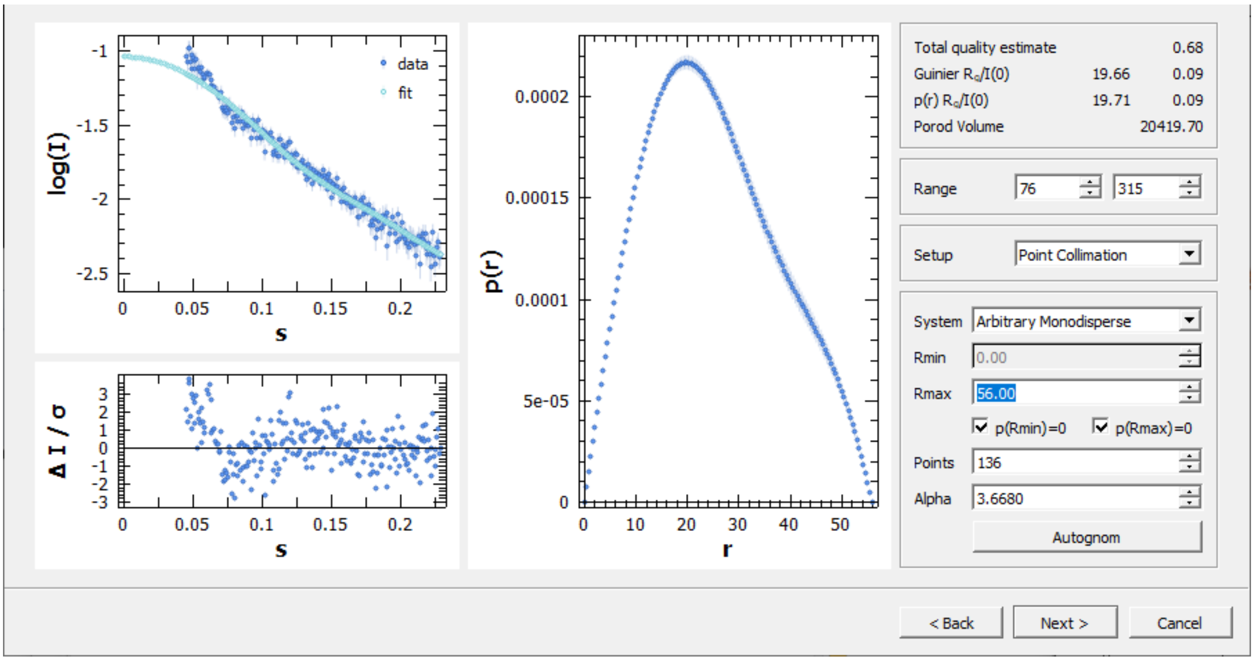


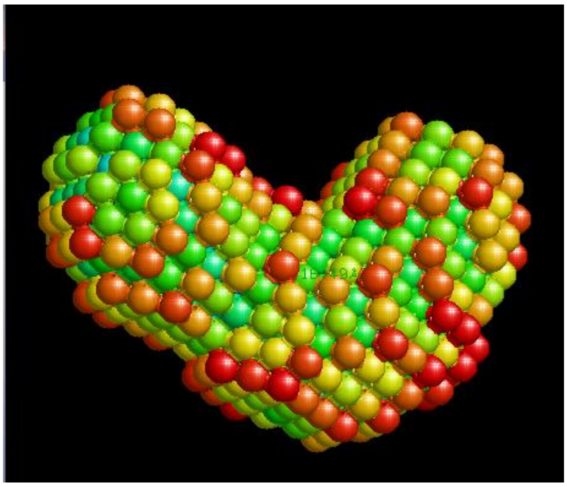

**Supplementary figure 16.** Approximation of SAXS data for p54_1 aptamer (upper graph) and modelling of aptamer shape (bottom graph).

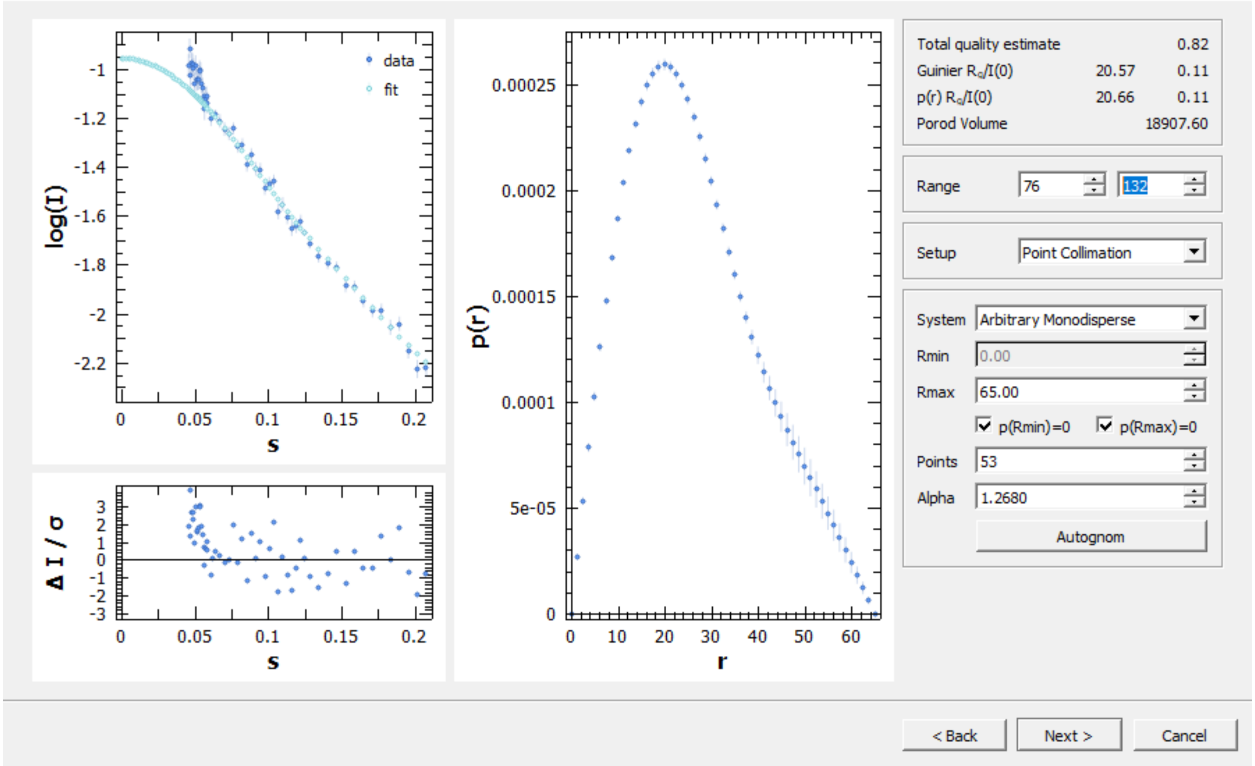


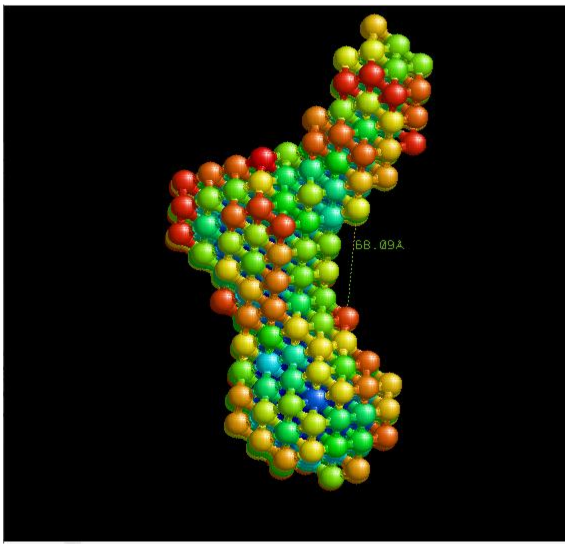


**Supplementary figure 17.** Approximation of SAXS data for p54_1 aptamer in the presence of $Ni^{2+}$ (upper graph) and modelling of aptamer shape (bottom graph).

**Supplementary table 4.** Lists of polar contacts in p54_1 aptamer and its complex with two nickel ions. 'WC bp' is a Watson-Crick base pair, 'h' means heterocycle, 'd' means deoxyribose, 'p' means phosphate. 'Hot spot' residue is a residue, which is involved in 3 or more polar contacts. The differences in polar contacts of two aptamer structures are shown in blue. The analysis was performed using PyMOL software.

| p54_1 | Secondary structure | p54_1 with $Ni^{2+}$ | Secondary structure |
|---|---|---|---|
| WC bp C5h-G55h | | WC bp C5h-G55h | |
| WC bp C6h-G54h | | WC bp C6h-G54h | |
| WC bp T7h-A53h | | WC bp T7h-A53h | |
| 1 contact C8h-G52h | | WC bp C8h-G52h | |
| 1 contact C8h-G18p | | 2 contacts G9h-T21p | |
| 1 contact G9h-G42h | | 1 contact G9h-C46h | |
| 1 contact G9h-A12h | | 2 contacts G10h-A12h | |
| 1 contact G10h-A12h | | 1 contact G10h-G39h | |
| 2 contacts G10h-G42h | | 1 contact G11s-C13h | |
| 1 contact G11h-G39s | | 1 contact A12h-G15h | |
| 1 contact G11h-A38s | | 1 contact A12h-G43h | |
| 2 contacts G11h-A38h | | 1 contact A12p-C13h | |
| 1 contact A14h-A41h | | 1 contact A14h-G43p | |
| 1 contact G15h-A41p | | 3 contacts G15h-G43h | |
| 2 contacts G15h-A41s | | 2 contacts G16h-A20h | |
| 1 contact G16h-G42h | | 1 contact G17h-G18s | |
| 1 contact G16h-G43h | | 1 contact G17h-C49h | |
| 2 contacts G17p-G52h | | 2 contacts G17h-A50h | |
| 2 contacts A20h-A48h | | 1 contact G18h-A50s | |
| 1 contact A20h-C49h | | 1 contact G18h-T51p | |
| 1 contact T21h-A35h | | 1 contact G18h-T51s | |
| 2 contacts T21h-A36h | | 1 contact T21h-C37h | |
| 1 contact T21s-G45h | | 1 contact T21h-G45p | |
| 1 contact A27h-G26h | | 1 contact G28h-T29h | |
| 1 contact A36h-G45p | | 1 contact A36h-G45p | |
| 1 contact C37h-C44h | | 1 contact G39h-C44s | |
| 1 contact C37p-C44h | | 3 contacts G39h-C46s | |
| 1 contact C37h-G45h | | 1 contact C40h-C44s | |
| 1 contact A38h-G45h | | 1 contact G43h-G45h | |
| 1 contact A38h-G43h | | 1 contact G43h-G45s | |
| 1 contact G39h-C40s | | 1 contact G43h-C46p | |
| 1 contact G39h-A41h | | | |
| 2 contacts G39h-G42h | | | |
| 1 contact G39h-G43h | | | |
| 1 contact G42h-G43h | | | |
| WC bp T22h-A35h | | WC bp T22h-A35h | |
| WC bp C23h-G34h | | WC bp C23h-G34h | |
| WC bp A24h-T33h | | WC bp A24h-T33h | |
| WC bp G25h-C32h | | WC bp G25h-C32h | |
| WC bp G26h-C30h | | WC bp G26h-C30h | |

### Materials and Methods

The inorganic salts and buffer components were purchased from AppliChem (Germany). The recombinant p54 protein with His tag was purchased from GeneTex (USA). HisLink protein purification particles from Promega (USA), T4 DNA ligase and Ultra II end repair / dA-tailing module from New England Biolabs (USA) were used. Magnetic particles AMPure XP beads, Blunt/TA ligase Master Mix, Native barcodes (NB01-24), Sequencing buffer and Library beads were obtained from Oxford Nanopore Technologies (UK). PCR kit was purchased from Syntol (Russia). All solutions were prepared using ultrapure water from the AquaLab facility (Russia).

#### OLIGONUCLEOTIDES

Oligonucleotides, including a randomized library, were synthesized by Lumiprobe LCC (Russia). The sequences of aptamers are shown in Supplementary table 2. Primers used for PCR of the aptamer-enriched library were 5'-GGTAGCATCGCTACGTC-3' and 5'-GATGCTACCCGTCATGT-3'; the complementary sequences are underlined.

#### BUFFERS

The composition of buffers used was as follows:

1) The selection buffer contained 100 mM HEPES pH 7.5, 100 mM NaCl, 50 mM KCl, 20 mM $CaCl_2$, 20 mM $MgCl_2$;
2) The selection buffer with $Ni^{2+}$ contained 100 mM HEPES pH 7.5, 100 mM NaCl, 50 mM KCl, 20 mM $CaCl_2$, 20 mM $MgCl_2$, 20 µM $NiCl_2$ (or with 20 mM $NiCl_2$ if specified in the text);
3) PBS contained 10 mM $Na_2HPO_4$, 1.8 mM $KH_2PO_4$ pH 7.3, 137 mM NaCl, 2.7 mM KCl;
4) Tris buffer with $Ni^{2+}$ contained 10 mM tris-HCl pH 7.0, 140 mM NaCl, 10 mM KCl, 20 µM $NiCl_2$.

#### VIRUSES

The study used the following strains of ASFV: Stavropol_01/08 (Genotype II, Serogroup 8, GenBank accession number PQ672299.1) [11], KK262 (Congo-a) (Genotype I, Serogroup 2, GenBank accession number OM249788.1) [12]. These strains were obtained from the collection of Federal Research Center for Virology and Microbiology. The Stavropol_01/08 strain was cultured in pig macrophage cells, while the KK262 strain was cultured in an inoculated COS-1 cell line. COS-1 cells were kindly provided by C. Gallardo from CISA-INIA, Spain. BA71V strain of ASFV (Genotype I, GenBank accession number NC_001659.2) [43] was obtained from the collection of Institute of Molecular Biology of NAS of (Armenia).

ASFV titer determination was performed using pig macrophage cell cultures with the addition of a 1% suspension of pig erythrocytes in 96-well plates for cell culturing. The infectious activity of the virus was determined by the Reed and Muench method [43] and expressed as lg $HAD_{50}$/mL. The titer was calculated based on three independent experiments. Inactivation of the virus for the binding experiments was achieved using β-propiolactone following the protocol described by Yang and Yang [45]. The titers were 7.00±0.19 lg $HAD_{50}$/mL for KK262 and 7.63±0.13 lg $HAD_{50}$/mL for Stavropol_01/08.

To estimate the viral titer, nanoparticle tracking analysis (NTA) was performed using the ZetaView PMX420-QUATT device (Particle Metrix GmbH, Germany). Data processing was performed with the ZetaView NTA software. Before starting measurements, the device was calibrated with a standard sample of polystyrene latex particles with a diameter of 100 nm with a known concentration (Applied Microspheres B.V., Netherlands), according to the manufacturer's instructions. Viral stock solutions were then diluted 100-fold with PBS. Particle concentration and size distributions were measured on 11 focal planes in 10 repeats. Camera sensitivity and exposure were individually adjusted for each series of experiments to achieve optimal signal-to-noise ratios. The virus titers were $1.3 \cdot 10^{10}$ particles/mL for KK262 cos-1 and $1.2 \cdot 10^{10}$ particles/mL for Stavropol 01/08 MF, and the virus diameters were 187 nm and 180 nm, respectively.

#### IMMOBILIZATION OF PROTEIN ON A SOLID CARRIER

The sorption capacity of HisLink particles was determined by measuring the reduction of optical density of in the protein solution after adsorption. Washing of the particles and adsorption of the protein were carried out in a buffer containing 100 mM HEPES pH 7.5 with 200 mM NaCl. After 60 minutes of protein immobilization, approx. 100 µg of protein was adsorbed onto 100 µL of particles, indicating a sorption capacity of 1 mg/mL for the particles.

### LIBRARY PREPARATION AND COUNTER-SELECTION CYCLE WITH HISLINK PARTICLES

The oligonucleotide library (740 nmol) was dissolved in 20 mL of the selection buffer to obtain a 37 µM solution. The solution was then heated for 10 minutes at 95$^{o}$C and cooled at room temperature to allow for the assembly of secondary DNA structures.

An aliquot of 100 µL of HisLink particles without protein was placed between two pieces of polyethylene terephthalate (PET) membrane with a pore diameter of 300 nm in a filter holder. The system was slowly washed, at about 1 mL/min, using the selection buffer (25 mL), with the OB1 MK3+ microfluidic system from Elveflow (France) equipped with an MFS flow sensor. After that, the oligonucleotide library was passed through the particles to separate oligonucleotides that could bind to off-target binding. Finally, the filter was rinsed with 15 mL of the selection buffer, and the oligonucleotide library that passed through the system was collected for further use in the positive selection cycle.

### SELECTION OF DNA APTAMERS FOR THE RECOMBINANT PROTEIN P54

A sample of 25 µg of recombinant p54 protein with a His-tag was immobilized on 100 µg of HisLink particles for 60 minutes with continuous stirring at 4°C in a buffer consisting of 100 mM HEPES pH 7.5 with 200 mM NaCl. The particles were then placed between two pieces of PET membrane with a pore size of 300 nm in a filter holder. The system was slowly rinsed with the selection buffer (15 mL) at a rate of 1 mL/min. Then, the oligonucleotide library, which was collected after the counter-selection cycle (volume of approx. 30 mL), was passed through the particles, and the system was rinsed with 15 mL of the selection buffer. The selection conditions were then gradually tightened, by increasing the detergent concentration step-by-step. First, 30 mL of the selection buffer with the addition of 0.01% Tween-20 was used, and then 30 mL with the addition of 0.1%. The fraction of oligonucleotides that remained on the HisLink particles after all washing steps was collected for further analysis.

### PURIFICATION OF THE DNA APTAMER LIBRARY

The DNA aptamer library was purified by treating the particles and membranes with 1 mL of Trizol for 5 minutes. Next, 200 µL of chloroform was added, and after a 3-minute incubation, the aqueous phase was separated by centrifugation (15 minutes at 12,000 g). The aqueous fraction was then divided into 100 µL aliquots in separate tubes. A volume of 1 mL of a 2% $LiClO_4$ in acetone was added to each aliquot, and the DNA was precipitated by freezing for 15 minutes at -20°C. After that, the precipitate was separated by centrifugation for 5 minutes at 12,000 g, and the procedure was repeated two more times to wash the precipitate with 1 mL of a 2% $LiClO_4$ in acetone. Finally, the precipitate was rinsed three more times with pure acetone and dried in air. The DNA was dissolved in 60 µL of ultrapure water, yielding a final concentration of 60 µM.

### PCR AMPLIFICATION OF DNA APTAMER LIBRARY

Primers were selected to complement both the library and each other, in order to increase the size of the PCR product, which is beneficial for the chosen sequencing method. A 25 µL PCR mixture was prepared, containing 10 µL of a 2.5x PCR reaction mixture, 1.5 µL of 25 mM $MgCl_2$, 11.5 µL of ultrapure water, 0.5 µL of 50 µM of each of the two PCR primers, and 1 µL of the matrix (3 µM). The PCR cycling conditions consisted of preheating at 95°C for 5 min, followed by 20 cycles of 95°C for 1 min, 30°C for 1 min, and 62°C for 1 min, and then 20 more cycles at 95°C for 1 min, 60°C for 1 min, and 62 °C for 1 min.

### PREPARATION OF THE LIBRARY FOR SEQUENCING

T4 DNA ligase was used to extend the library. For this purpose, the PCR mixture was mixed with a buffer containing 66 mM tris-HCl at pH 7.6, 10 mM $MgCl_2$, 1 mM ATP, 1 mM dithiothreitol, and 7.5% PEG of molecular weight 6000 Da. 65 µL (1/10 of the total volume) of T4 DNA ligase with an activity of 400 U/µL was then added. The mixture was incubated overnight at 20$^{o}$C. After ligation, the reaction product was purified using acetone and $LiClO_4$, following the protocol described above.

The ligation product was processed with a set of Ultra II end repair/dA-tailing module to prepare the library ends for barcode ligation. For this purpose, three test tubes were prepared, each of which was filled with 5 µL of the ligation product, 7.5 µL of water, 1.75 µL of Ultra II end-prep reaction and 0.75 µL of Ultra II end-prep enzyme mix. The mixture was incubated for 20 minutes at room temperature, followed by inactivation of the enzyme by heating at 65°C for 5 minutes. Each DNA sample was then purified using magnetic particles, AMPure XP beads. 15 µL of magnetic particles were mixed with the DNA for 5 minutes, after which the supernatant was removed, and the magnetic beads were washed twice with 200 µL of 80% ethanol. Finally, 10 µL of water, 2.5 µL of Native barcodes barcoding sequence (SQK-NBD114-24), and 10 µL of Blunt/TA ligase Master Mix were added. The mixture was incubated overnight at +4°C.

### SEQUENCING OF THE LIBRARY

The libraries of DNA aptamers from three test tubes were mixed together. 12 µL of the mixture was combined with 37.5 µL of the Sequencing buffer and 25.5 µL of the Library beads. This mixture was then added to a FLO-MIN114 cell of a MinION device (Oxford Nanopore Technologies, UK). Sequencing continued until the cell had completely lost its activity (approx. 72 hours). Basecalling was performed using a High-accuracy model 400bps with MinKNOW software (Oxford Nanopore Technologies, UK). As a result, approx. 70 to 79% of the reads were selected by the software as having passed quality control with a minimum Q score of 9.

### EVALUATION OF THE AFFINITY OF APTAMERS BY BIOLAYER INTERFEROMETRY

The affinity was evaluated in several ways. In the first approach, the recombinant p54 protein was immobilized on the surface of an amine reactive 2nd generation sensor (Sartorius, Germany). The biosensors were hydrated for 10 min in water. The biosensors were then activated by 200 mM carboxymethylethanolamine and 100 mM sodium salt of N-hydroxysulfosuccinimide for 5 min. The p54 protein was diluted to a concentration of 0.16 mg/mL in 40 mM phosphate buffer with a pH of 6.4. The sensor was then washed and blocked with 100 mM HEPES buffer with a pH of 7.5 for 3 min. After that, the signal was stabilized in a buffer (PBS or the selection buffer with varying concentrations of $Ni^{2+}$). Potential aptamers without modification were folded in the same buffer. Specifically, a 1 µM solution of the aptamer was heated for 5 minutes at 95°C, and then cooled at room temperature. The biosensors with immobilized protein interacted with 1 µM, 500 nM, 250 nM, and 125 nM solutions of the aptamers. The curves of association and dissociation of the complex were obtained using the Bli instrument (Forte-Bio, USA). The constants were calculated using BliMart software developed by Pavel Martynyuk. The software approximated the dissociation curves with an exponential function $y=y_0+A\ exp(-Bx)$ with $y_0$=0. The exponential factor $B$ corresponds to the dissociation rate constant $k_d$. The association curves were also approximated with the same exponential function with $B=k_d+k_a\ C_{apt}$, where $C_{apt}$ is the aptamer concentration and $k_a$ is the association rate constant. Finally, the equilibrium dissociation constant of the complexes were calculated using the equation $K_D=k_d/k_a$.

To assess the affinity of aptamers to ASFV virions, biotinylated DNA aptamers were folded in the buffer and immobilized on the surface of streptavidin sensors (Sartorius, Germany). Virus stock solutions were then diluted 5-, 20-, and 50-fold with PBS. Non-specific binding was subtracted from the curves obtained with a non-specific aptamer, namely an aptamer for the mycotoxin T-2 (biotin-5'-GTATATCAAGCATCGCGTGTTTACACATGCGAGAGGTGAA-3' [46]). The constants were calculated using BliMart software developed by Pavel Martynyuk.

### EVALUATION OF VIRUS BINDING TO APTAMERS BY FLUORESCENCE POLARIZATION ASSAY

The FAM-labeled aptamers were folded in PBS according to the protocol described above. Aliquots of 5 µL of ASFV stock solutions were added sequentially to 1 mL of 3-5 nM aptamer solutions in PBS. After a 1-minute incubation, fluorescence polarization (mP) was measured three times using the SENTRY 200 instrument (Ellie, USA). The non-specific binding of the non-specific sequence p54_8_FAM was subtracted from the experimental curves to obtain the specific binding. The $K_D$ was calculated using OriginPro software (OriginLab, USA) with the Hill equation: $f_b = (B_{max} \cdot C_{ASFV}^{n})/(K_D^{n} + C_{ASFV}^{n})$, where $f_b$ is the fraction of bound aptamer, $B_{max}$ is the maximum signal, and $C_{ASFV}$ is the virus concentration, and $n$ is the Hill coefficient.

### EVALUATION OF VIRUS BINDING TO APTAMERS BY NANOPARTICLE TRACKING ANALYSIS

Alexa488-labeled aptamers were folded into PBS according to the protocol described above. 20 µL of ASFV stock solution was added to 20 µL of 2 µM aptamer solution. After 15 minutes of incubation, 500 µL of PBS was added and filtered through a membrane with a pore diameter of 0.06 µm to filter out the unbound aptamers. Then, virus particles were dissolved in 2 mL of PBS, and the resulting suspension was analyzed by the NTA on a ZetaView PMX420-QUATT device (Particle Metrix GmbH, Germany); data processing was carried out using the ZetaView NTA software. Before starting the measurements, the device was calibrated using a standard sample of polystyrene latex particles labeled with Alexa-488 (Applied Microspheres B.V., the Netherlands), following the manufacturer's instructions. The particle concentration and size distribution were measured on 11 focal planes in 5 rotations. The sensitivity and exposure settings of the camera were selected individually for each series of experiments to achieve an optimal signal-to-noise ratio in the light scattering and fluorescence modes. The number of labeled particles was divided by the total number of particles, resulting in a fraction of bound viruses.

**ESTIMATION OF ANTIVIRAL EFFECTS OF APTAMERS**

Vero 76 cells were seeded in a 48-well plate. Growth medium was minimum essential medium (MEM) from Biowest (France) with 10% fetal bovine serum (FBS) from Capricorn (Germany). The cells were incubated for 24 hours in 5% $CO_2$ and 37$^{o}$C. After incubation, the growth medium was aspirated.

The co-treatment protocol:

Cells were washed with warm PBS to completely remove residual growth medium. Then, 10 µM aptamer solutions were added to the cells and incubated for 1 hour in the incubator. Then Vero-adapted BA71V strain of ASFV of Moi 0.2 was added directly to the medium and incubated for 1 hour in the incubator (co-incubation of cells, substances, and virus). Afterwards, the medium containing the virus and aptamers was aspirated. Cells were washed with PBS. Then, growth medium (MEM + 2% FBS) with 10 µM of aptamer was added. Incubation was 72 and 96 hours. After the incubation, medium from all wells was collected and analyzed by Virus yield reduction $TCID_{50}$ assay.

The post-infection protocol:

Cells were washed with warm PBS to completely remove residual growth medium. Then Vero-adapted BA71V strain of ASFV of Moi 0.2 was added directly to the medium and incubated for 1 hour in the incubator (incubation of cells and virus). Afterwards, the medium containing the virus was aspirated. Cells were washed with PBS. Then, growth medium (MEM + 2% FBS) was added; the incubation was 16 hours. Next, the growth medium was aspirated, and new growth medium (MEM + 2% FBS) with 10 µM of aptamer was added. Incubation was 120 hours. After the incubation, medium from all wells was collected and analyzed by Virus yield reduction $TCID_{50}$ assay.

**CIRCULAR DICHROISM SPECTROSCOPY AND UV SPECTROSCOPY**

Aptamer solutions were folded at a concentration of 1 µM in PBS or the selection buffer with varying $Ni^{2+}$ concentrations. The solutions were placed into quartz cuvettes with an optical path length of 1 cm. Circular dichroism (CD) and UV spectra were acquired using a Chirascan spectrometer (Applied Photophysics, UK) and a MOS-500 spectrometer (BioLogic, France), equipped with a thermoelectric temperature controller. Spectra were acquired in the wavelength range of 240-360 nm, with the buffer spectrum serving as the baseline. Samples were heated at an average rate of 1.0°C/min. Melting experiments were conducted over a temperature range of 10-85°C. Melting temperatures were determined based on the dependence of molar CD intensity at 270 nm and UV absorbance at 260 nm on temperature. The curves were approximated using the Boltzmann function $y=A_2+(A_1–A_2)/(1+\exp((x-x_0)/dx))$, where parameter $x_0$ is the melting temperature. The parameters were determined using OriginPro software (OriginLab, USA).

**STUDY OF COMPLEX FORMATION BETWEEN APTAMERS AND $NI^{2+}$**

Stopped flow kinetics experiments were carried out using a MOS-200 rapid kinetics optical system equipped with an SFM-3000/S stopped-flow mixer (BioLogic Science Instruments, Seyssinet-Pariset, France). FAM-labeled aptamers were folded in the selection buffer with a final concentration of 1 µM. $NiCl_2$ solutions were added to the selection buffer at concentrations of 10 µM or 100 µM. Aptamers and cations were mixed in different volume ratios (1:1, 2:1, 3:1, 4:1, and 5:1), and fluorescence changes were measured light over a period of 20 seconds with excitation at 495 nm and a filter of 530 nm for the emitted. Each ratio was repeated 5-7 times, and the average curve was plotted. The data were approximated by an exponential function $y=y_0+A\ exp(-k_{obs}\cdot x)$ using OriginPro software (OriginLab, USA). The observed rate constant, $k_{obs}$, was plotted against $C_{Apt}\cdot C_{Me2+}^n$ in order to find the $n$ value, which provided a linear dependence. The intercept of this linear dependence with the ordinate axis corresponds to $k_{diss}$; whereas the tangent of the tilt angle corresponds to $k_{ass}$. The equilibrium dissociation constants of for the complexes were calculated using the equation $K_D=k_{diss}/k_{ass}$. Additionally, a model of two unequal sites was used, approximating with two exponential dependencies. In this case, each $k_{obs}$ was calculated separately.

### MOLECULAR MODELLING OF APTAMER STRUCTURE

Aptamer models were generated using AlphaFold 3. Due to the inability to generate aptamer structures containing nickel, cobalt and copper—elements adjacent to nickel in the periodic table—were used for this purpose. In all models except model 5, nickel ions were manually placed into cavities that were large enough to accommodate them and featured a high concentration of negative charge along their boundaries. These sites were usually observed near the bulged out nucleotides and bendings of the sugar-phosphate backbone. The ionic bonds in such contacts were mainly with oxides of phosphate groups. Less frequently, such sites were observed in narrow inter-chain regions and in some loops, where the ions formed bonds with the negatively charged atoms of the aromatic rings of the nucleotide bases. The model containing a hybrid G-quadruplex and a supporting tetrad was constructed using the strategy described by V.B. Tsvetkov [40]. The selection buffer was imitated by adding all buffer-related cations in the appropriate stoichiometries around the aptamers. All models were energy-minimized according to the protocol described in [40]. The models were verified by molecular dynamics simulations during 80 ns using Amber 22 software. The structural stability of the models and their free energies within the generalized Born approach were evaluated as described in the work already mentioned [40].

### NMR SPECTROSCOPY

For NMR titration experiments, the aptamer p54_1 was prepared at a concentration of 0.3 mM in a buffer containing 10% (v/v) $D_2O$. Aliquots of 50 mM $Zn^{2+}$ or $Ni^{2+}$ stock solutions were added to the aptamer samples in Shigemi NMR tubes to achieve aptamer-to-ion molar ratios of 1:0.5, 1:1, 1:2, and 1:5. $^1$H NMR spectra were acquired using a Bruker Avance 600 MHz spectrometer equipped with a room-temperature probe and a Bruker Avance III 700 MHz spectrometer equipped with a Prodigy triple-resonance cryoprobe. The measurement temperature was optimized between 293 and 298 K, and all subsequent experiments were performed at 298 K. Data were acquired using the zgpw5 pulse program with WATERGATE water suppression. Spectra were processed and analyzed using TopSpin software.

### SMALL-ANGLE X-RAY SCATTERING (SAXS) MEASUREMENTS

Small-angle X-ray scattering (SAXS) data were collected using a Xeuss 3.0 laboratory scattering system (Xenocs, France) equipped with a high-brightness GeniX3D Cu $K_\alpha$ microfocus X-ray source ($\lambda$ = 1.5418 Å) operated at 50 kV and 0.6 mA. The collimation system was adjusted to the high-resolution beam mode utilizing two motorized scatterless slits: S1 (0.8 x 0.8 mm$^2$) and S2 (0.5 x 0.5 mm$^2$). The scattering patterns were recorded using a hybrid photon-counting 2D detector Eiger2 R 1M (Dectris, Switzerland). To cover a comprehensive range of the momentum transfer vector Q defined as $Q = 4\pi \sin\theta/\lambda$, where $2\theta$ is a scattering angle, a dual-distance configuration strategy was employed. Measurements for medium-angle regions (MAXS topology) were carried out at a sample-to-detector distance of 400 mm, while low-angle regions (SAXS, Guinier domain) were recorded at a sample-to-detector distance of 1800 mm.

The liquid samples of DNA aptamers (5 mg/mL) and their corresponding background buffers were loaded into 1.0 mm borosilicate glass capillaries. Transmittance and sample alignment were automatically verified prior to each run via automated X-scan profiling. To optimize the signal-to-noise ratio and eliminate potential inter-molecular radiation damage artifacts, automated frame-by-frame data collection was implemented. Both buffers and DNA aptamer solutions were exposed for 240 minutes per sample, accumulated as a series of 12 independent 20-minute frames. All measurements were performed at a room temperature (~22°C) under vacuum.

### STRUCTURAL PARAMETER ESTIMATION AND *AB INITIO* MODELING

The data analysis, structural parameter extraction, and three-dimensional shape reconstruction were carried out using the standard modules of the ATSAS 3.0 software package. The radius of gyration ($R_g$)

and scattering intensity (I(0) were determined via Guinier analysis using the PRIMUS wizard within the linear region satisfying the condition ($Q_{max} \cdot R_g < 1.3$).

The pair-distance distribution functions, P(r), and the corresponding maximum particle dimensions ($D_{max}$) were evaluated by indirect Fourier transform using the GNOM program. The total excluded molecular volume (Porod volume, $V_p$) was calculated from the smoothed scattering curves to estimate the apparent molecular mass (Mw) assuming a standard empirical density factor of Mw ≈ Vp / 1200.

Three-dimensional ab initio low-resolution bead models were reconstructed using the DAMMIN algorithms. For each configuration, 20 independent simulation runs were executed in slow mode assuming no particle symmetry (P1).
